# Application and Characterization of the Multiple Instance Learning Framework in Flow Cytometry

**DOI:** 10.1101/2025.06.10.658646

**Authors:** Zhiyuan Ding, Alexander Baras

## Abstract

For decades, flow cytometry has allowed for single-cell profiling based on selected biomarkers and is widely used in both clinical and research settings. One major limitation of most conventional flow cytometry analyses is the dependency on a mostly manual gating process. This generally involves sequentially selecting biomarkers to isolate phenotype-associated cell populations, an approach that is both labor-intensive and prone to bias. To address this challenge, we introduce the application of a series of multi-instance learning frameworks for automated flow cytometry data analysis. Our models demonstrate strong performance across diverse biomedical applications, including cancer subtyping based on tumor-infiltrating immune cells, HIV survival stratification, AML minimal residual disease prediction, and COVID-19 severity assessment. We further examine how network architecture affects predictive performance and the detection of rare but clinically significant cell populations. Notably, our models utilize attention mechanisms to directly identify phenotype-associated cell subsets, serving as an interpretable, data-driven alternative to fully manual gating. These findings underscore the potential of multi-instance learning as a scalable and interpretable framework for flow cytometry, with broad applications in precision medicine and translational immunology.

## Introduction

Conventional flow cytometry (FCM) is an advanced single-cell technology that uses a variety of detection modalities to characterize sets of biomarkers per cell. Modern advancements like spectral fluorescence and mass cytometry have enabled the characterization of complex sets of biomarkers with single-cell resolution, allowing for the identification of cell populations with over 40 parameters [1]. Over the past 30 years, flow cytometry has been widely applied in various fields, including immunophenotyping and abnormal cell population identification clinically. The conventional analytic approach involves demarcating cell populations based on predefined biomarker value ranges (commonly referred to as “gating strategy”) and applying these selections to other markers within the experiment for further study. However, this approach is somewhat subjective, labor-intensive, and becomes increasingly limited as the number of biomarkers grows beyond approximately 5–10. As a result, relationships among tens or even hundreds of markers may go undetected, increasing the risk of missing biologically relevant cell populations that may help better describe the biological phenomenon being studied.

Of course, a variety of computational approaches have been previously utilized with FCM data, initially relying on conventional machine learning approaches. These methods, ranging from early approaches such as FLOCK [2], SPADE [3], and Citrus [4], to more recent ones such as XShift [5], FlowSOM [6], VaPo [7], DeepCyTOF [8], PhenoGraph [9], and FAUST [10], typically follow a stepwise process to identify cell populations and then use this information for specific tasks, such as detecting disease-associated populations. For a comparison of these strategies, readers may refer to [11] and [12], which provide benchmarking analyses. However, these approaches have several limitations [13], including information loss during the cell gating process, sensitivity to the selected clustering method, and an inability to detect cellular changes that are not linked to well-defined populations.

As an alternative approach, some deep learning-based methods have begun to be employed in FCM analysis due to their ability to perform comprehensive cellular analysis without relying on traditional cell gating or clustering, which could be considered a form of “hand-crafted feature engineering”. The first neural network based approaches focused on extracting cellular features across the biomarkers that were measured in FCM data, utilizing carefully designed network architectures to model cytometry information. For instance, CellCNN [14] employs a single 1D convolutional layer to identify disease-associated cellular features, while [13] leverages multiple convolutional layers for “deeper” cellular feature extraction, akin to the concept of deep learning derived features. Similarly, CytoSet [15] utilizes a multi-layer perceptron (MLP) to enhance the feature extraction process. However, these approaches have several limitations. First, they are heavily focused on extracting meaningful cellular features without selectively utilizing specific cells for prediction. Second, the interpretability of these frameworks is often unclear or requires extensive processing steps, such as directly using activation values from convolutional layers [14]. Finally, most methods incorporate a subsampling process, typically limiting the number of analyzed cells per sample to a fixed count, often no more than 10,000, which may not fully capture cellular heterogeneity.

To address these limitations, this work employs multi-instance learning (MIL) [16, 17] for FCM analysis, where individual cells are treated as instances, and disease-associated cells are selectively identified using cellular-level attention mechanisms optimized through sample-level prediction. Previous works that use MIL in FCM analysis for leukemia include [18–20]. In [18], attention-based MIL (ABMIL) [17] is applied for AML subtyping across different cohorts of AML tasks. [19] presents a benchmarking study of various computational models for FCM analysis using both in-house and public AML datasets. In [20], a Cell Scoring Neural Network or vanilla MIL is employed for leukemia identification. However, previous works have several limitations. First, they all assume a single selected set of cells for the downstream sample-level task, which fails to fully capture the heterogeneous landscape of cell populations. Additionally, the associations between selected cells and phenotypes (sample labels) cannot be directly identified. Second, existing studies primarily focus on hematological analysis, while other important FCM applications, such as cancer subtyping, survival analysis, and immunoreactive analysis, remain underexplored. Finally, previous research predominantly relies on a single selected architecture, without fully examining differences across various MIL architectures. Nevertheless, in FCM analysis, where sample sizes are relatively small, adopting more complex network structures does not necessarily lead to better performance.

In this work, we address these limitations by benchmarking MIL structures across various FCM tasks, evaluating both sample-level prediction performance and cellular-level interpretability. Specifically, we design vanilla MIL and ABMIL with interpretable attention heads linked to sample phenotypes and assess their performance across different scenarios, including cancer subtyping based on the FCM of the associated immune cells [21], survival analysis in the context of HIV infection /exposure [22], minimal residual disease (MRD) analysis in leukemia [23], and immune profiling for COVID-19 severity [24]. The results demonstrate that the proposed MIL frameworks not only achieve accurate sample-level predictions but also effectively identify phenotype-associated sensitive cell populations across different tasks.

## Results

### Interpretable MIL framework design

We design MIL frameworks, including vanilla MIL (vMIL) and ABMIL, specifically tailored for FCM data (see Architecture details for details, Fig. 1). Both frameworks take raw FCM data as input. In this context, cells from the biological domain are treated as instances in the MIL scenario.

**Figure 1.**
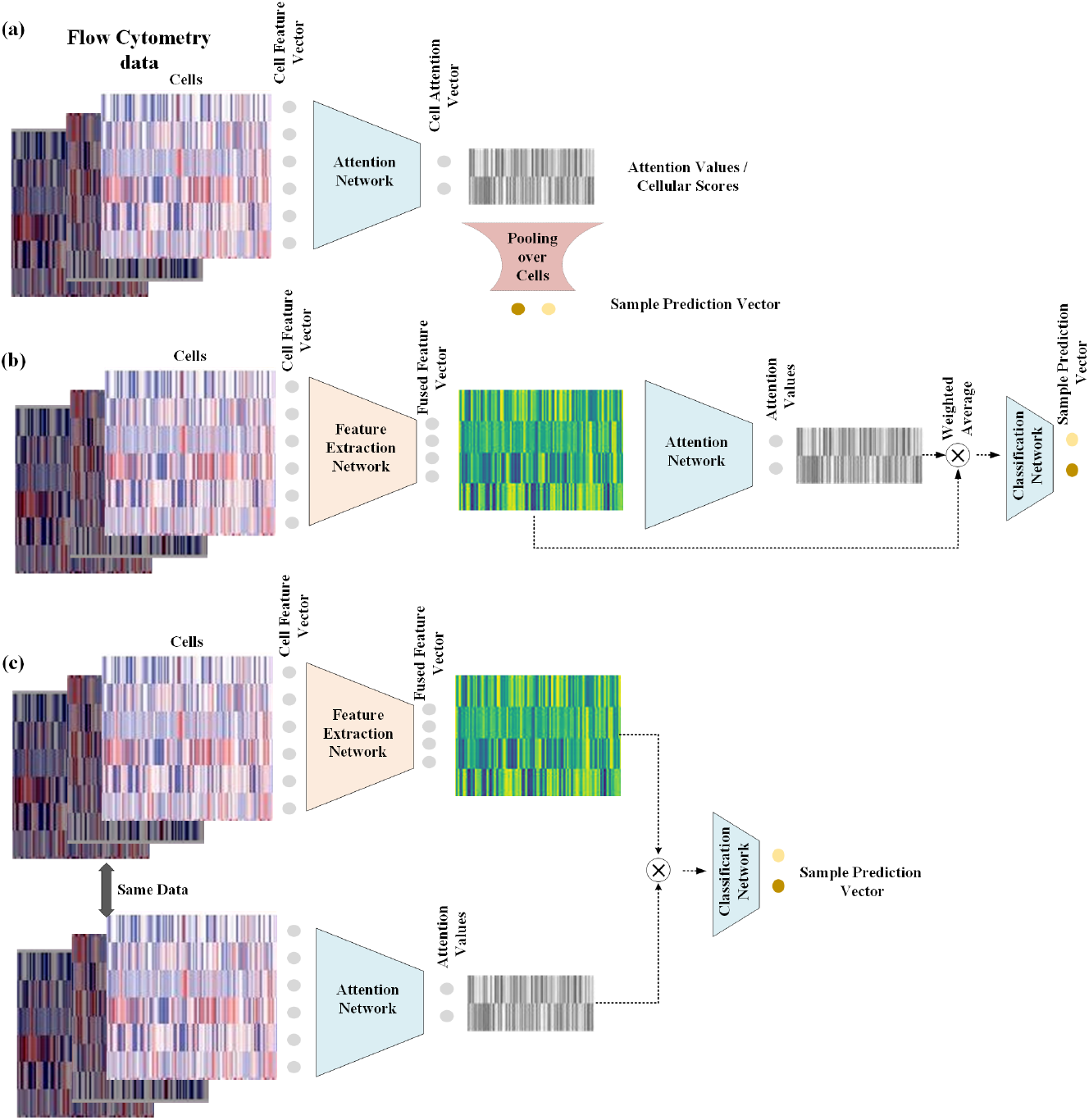
MIL frameworks compared in this paper. (a) **vMIL.** (b) Dependent ABMIL (**dep-ABMIL**), where the instance attention network receives input from the instance feature extraction network, creating a dependency between the two. (c) Independent ABMIL (**ind-ABMIL**), where the instance attention network operates independently from the instance feature extraction network, taking input directly from the raw features.

In vMIL, cellular associations with each class are extracted using an instance-level (cellular) attention MLP and then aggregated at the bag (sample) level for prediction. By incorporating a well-designed instance-level attention mechanism and bag-level aggregation strategy, vMIL effectively identifies cell populations associated with each phenotype.

We design ABMIL following the pipeline from [17], where, unlike vMIL, cells are selected through an additional attention network. Instance latent variables, extracted via a feature-extracting MLP, are then averaged but weighted by the learned attention to generate bag-level representations for downstream tasks. To ensure interpretable cellular attention, we design the sample-level MLPs to be class-specific and combine them only once at the final bag-level output. Rather than using gated attention, we employ a simple MLP to represent the instance attention network. Additionally, we explore two ABMIL variants for meaningful instance score generation, differing in the degree of entanglement between the cellular feature extraction network and the attention network (Fig. 1). Unless otherwise specified, the number of attention heads is set equal to the number of sample classes.

In the following results, we identify key settings impacting prediction performance and model interpretability. These include instance attention activation function, which reflects the assumed probability distribution of cellular attention (see Supplementary Instance activations distribution assumption.), as well as attention aggregation and normalization strategies. For each of these critical settings, we conduct an extensive search over their combinations while also sampling other hyperparameters, including training configurations and network architectures (see Experiment settings), to identify the structure that achieves the best predictive performance. All presented results are based on 5-fold cross-validation.

### MIL captures phenotypic distinctions in human tumor infiltrates

We use the CD8+ tumor-infiltrating lymphocytes (TIL) dataset [21], sampled from colorectal and lung tumors, to demonstrate MIL’s ability to identify phenotype-specific cell populations. After preprocessing (details in Dataset details and preprocessing), the TIL dataset consists of 76 samples, including 47 cases of colorectal cancer (CLC) and 29 cases of lung cancer (LC). In the TIL dataset, the number of cells per sample ranges from fewer than 200 to over 200, 000, with each cell characterized by 22 biomarkers.

MIL frameworks are able to achieve high accuracy (accuracy 0.9057, area under the receiver operating curve (AUC) 0.9454, macro F1 score 0.9075) when using suitable instance activation and bag normalization strategies (Fig. 2 (a)). We also observed that a more complex network structure (ABMIL compared to vMIL) does not necessarily improve sample-level prediction performance in the TIL dataset (*P* = 0.0842). The bag normalization method significantly impacts sample-level prediction performance (*p* = 0.0077), while the instance activation function shows no statistically significant effect on sample prediction (*p* = 0.9665). Moreover, when extending our observations to all experimental trials with varying hyperparameter settings (e.g., learning rate, dropout, instance feature extraction MLP, and instance attention MLP, see Supplementary Fig. 1 (a)), the impact of network structure (*p* = 4.47 *×* 10^−31^) and bag normalization (*p* = 3.82 *×* 10^−16^) on performance becomes significant, while instance activation remains insignificant (*p* = 0.3809). Performance is also compared against existing cellular clustering approaches. Following [11], top-performing methods, including PACMAN [25], FlowClust [26], FlowSOM [6], and Depeche [27], were selected. The results (Supplement Table 1) indicate that although MIL frameworks do not require additional cellular clustering steps, their performance remains comparable to established clustering-based approaches.

**Figure 2.**
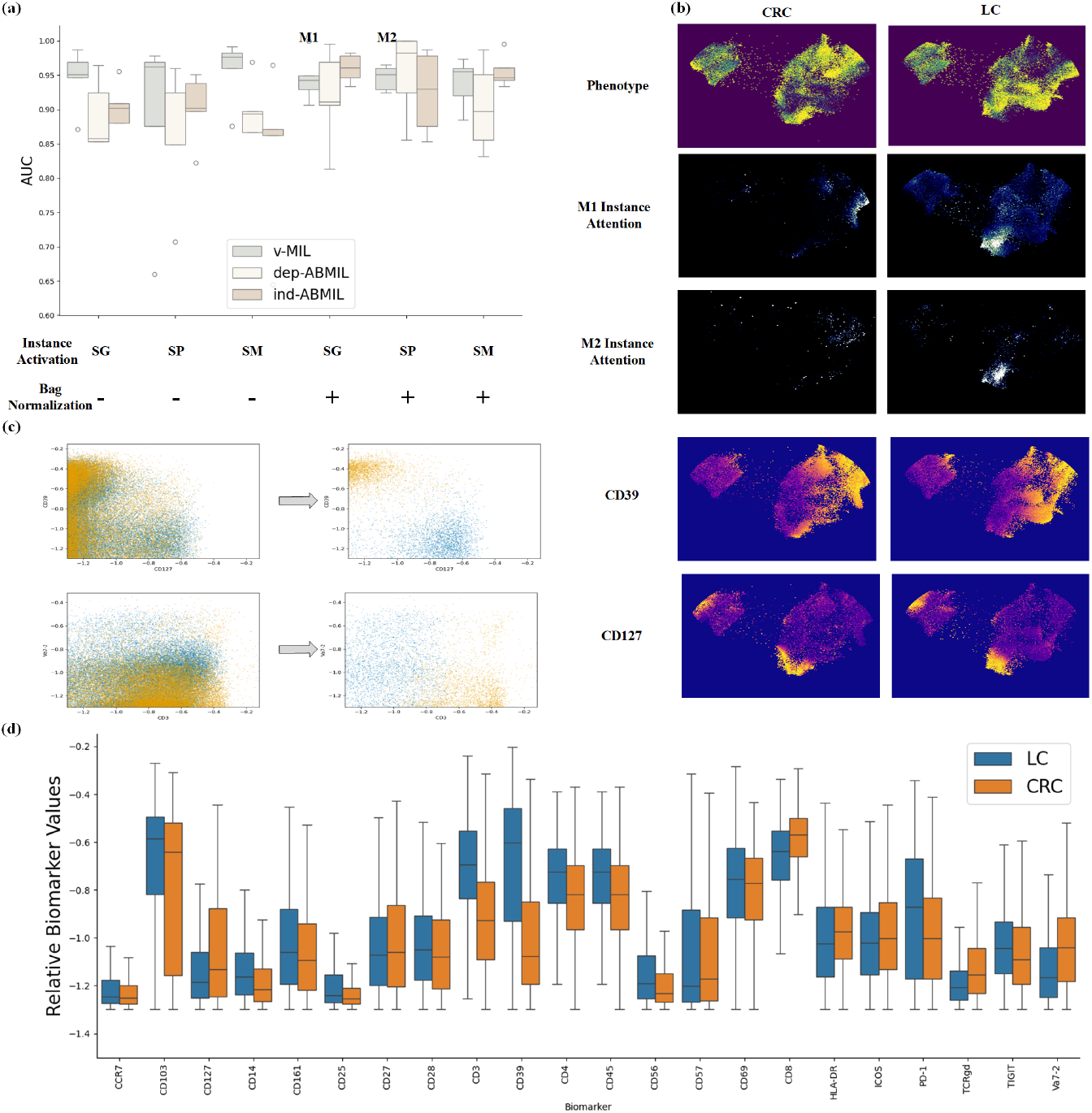
(a) Best CRC and LC subtyping performance across different MIL framework settings, including MIL structures, instance activation, and bag normalization strategies. The evaluated instance activation methods include Sigmoid (SG), Softplus (SP), and Softmax (SM). Bag normalization methods across attention heads are combined with specific aggregation strategies: averaging across cells without bag normalization (−) and summing across cells with bag normalization (+). Models M1 and M2 are used for subsequent interpretative analysis. (b) UMAP visualizations of 60,000 randomly sampled cells under different conditions. The first row shows cell distributions from CRC and LC samples. The second and third rows illustrate class-specific attention maps from models M1 and M2, respectively. The fourth and fifth rows depict CRC- and LC-sampled cell activation responses for biomarkers CD39 and CD127, selected based on (d). (c) Simulation of gating results based on model attention. The left panel displays cell distribution before filtering, while the right panel shows cell distribution after filtering. CRC (orange) and LC (blue) cells are color-coded accordingly. (d) Biomarker values are statistically analyzed for cell populations with high LC and CRC attention values (top 5%) of M1, derived from CRC and LC sampled cells. CD127, CD3, CD39, and =v*α*7.2 are identified as discriminative biomarkers and used for further analysis.

Trained MIL frameworks provide interpretable instance-level attention and identify cell populations associated with specific phenotypes. Phenotype-associated cell populations exhibit a more distinct biomarker distribution than the unfiltered population (*p* = 0.0081, Fig. 2 (b)). We project the biomarker values into UMAP [28] space and visualize the given instance attentions. Despite the heterogeneous relationship between phenotype and cell populations (Fig. 2 (b) 1^*st*^ row), MIL models effectively identify tumor-associated cell populations (Fig. 2 (b) 2^*nd*^ and 2^*rd*^ rows), demonstrating consistency across varied settings. The identified CRC- and LC-associated cell populations exhibit strong associations with several biomarkers (Fig. 2 (d)), including CD39, CD127, CD3, and v*α*7.2 (*p* ≪ 0.001, visualization of all 22 biomarkers in Supplementary Fig. 2). According to the trained model M1, LC-associated cell populations are characterized by a CD39+CD127-CD3+v*α*7.2-profile, whereas CRC-associated cell populations display a CD39-CD127+CD3-v*α*7.2+ property relatively [29, 30]. Similar to the manual gating process in FCM analysis, we visualize instance attention in 2D space before and after model (M1) selection using selected channels (Fig. 2 (c)). The results indicate that the models enable the selection of distinctive cell populations from CRC and LC, just as the gating process.

Consistency across different model settings and cross-validation ensures a stable and reliable analytical process. Cell populations are heterogeneous for a given model setting (Supplementary Fig. 3) and sample-level prediction performance varies due to differences in dataset distribution across splits (Fig. 2 (a)). However, with appropriate network configurations, the identified cellular populations consistently follow the same pattern across dataset splits (e.g. in M1 Spearman Correlation *ρ*_*s*_ = 0.8900, Supplementary Fig. 4). Although instance activation strategies do not significantly impact sample-level prediction performance, the identified important cell populations may either follow a consistent pattern (M1 vs M2, Spearman Correlation *ρ*_*s*_ = 0.9197, Supplementary Fig. 4) or vary across different activation strategies while maintaining the same model structure (Supplementary Fig. 4 and 5)). The identified cell populations exhibit more complex patterns with different model architectures, such as vMIL compared to ABMIL. Notably, we observe that models can learn ‘opposite attention’ to optimize sample-level predictions, meaning the model may aggregate CRC-associated cell populations to compute LC-related representations and vice versa (Supplementary Fig. 5 M5). To address this issue, we propose using the learned vMIL as pretrained weights for initializing ABMIL’s instance attention MLP. This strategy, specifically designed for ABMIL with independent attention and feature extraction networks, effectively mitigates the observed issues (Supplementary Fig. 5 M6). Analysis of identified cell populations across different samples reveals that, despite variations in instance distribution, the identified CRC- and LC-associated cell populations remain consistent (Supplementary Fig. 3). When using Softmax activation, setting the number of attention heads equal to the number of phenotypes can be overly restrictive, as it forces each instance to be associated with exactly one phenotype. To address this limitation, we introduce an additional, unassigned attention head that is not linked to any specific phenotype, allowing the model greater flexibility in capturing ambiguous or non-informative instances (Supplementary Fig. 7). We also find that introducing multiple attention heads does not lead to better performance and more interpretable results (Supplementary Fig. 6).

### MIL achieves survival analysis on HIV

HIV natural history (HIVNH) dataset [22] benchmark MILs performance for survival analysis. We first reframe the survival prediction as a classification problem, predicting whether a sample corresponds to a patient who dies within four years or survives for at least five years. Based on this stratification, the final dataset consists of 87 patients who died within four years (poor outcomes) and 60 samples from those who survived at least five years (good outcomes). The HIVNH dataset presents a challenge due to its large cell size (up to one million) relative to its smaller sample size and limited number of biomarkers (10 channels).

Given the substantial computational burden of hyperparameter searching on the HIVNH dataset, we select the best-performing frameworks (vMIL, ind-ABMIL, and dep-ABMIL) from the TIL dataset and train them on the HIVNH dataset for verification. vMIL achieves slightly better performance than the ABMIL frameworks, with an accuracy of 0.6609, AUC of 0.7275, and a macro F1 score of 0.6871 (Fig. 3 (a)). However, the performance differences between models are not statistically significant (*p* = 0.4271). To further assess whether the model learns a meaningful prediction strategy, we stratify samples into two groups using vMIL predicted scores, trained on binary classification. The stratification results (log-rank test *p* = 0.0009, Fig. 3 (b)) indicate that the predicted scores effectively distinguish the dataset into subgroups.

**Figure 3.**
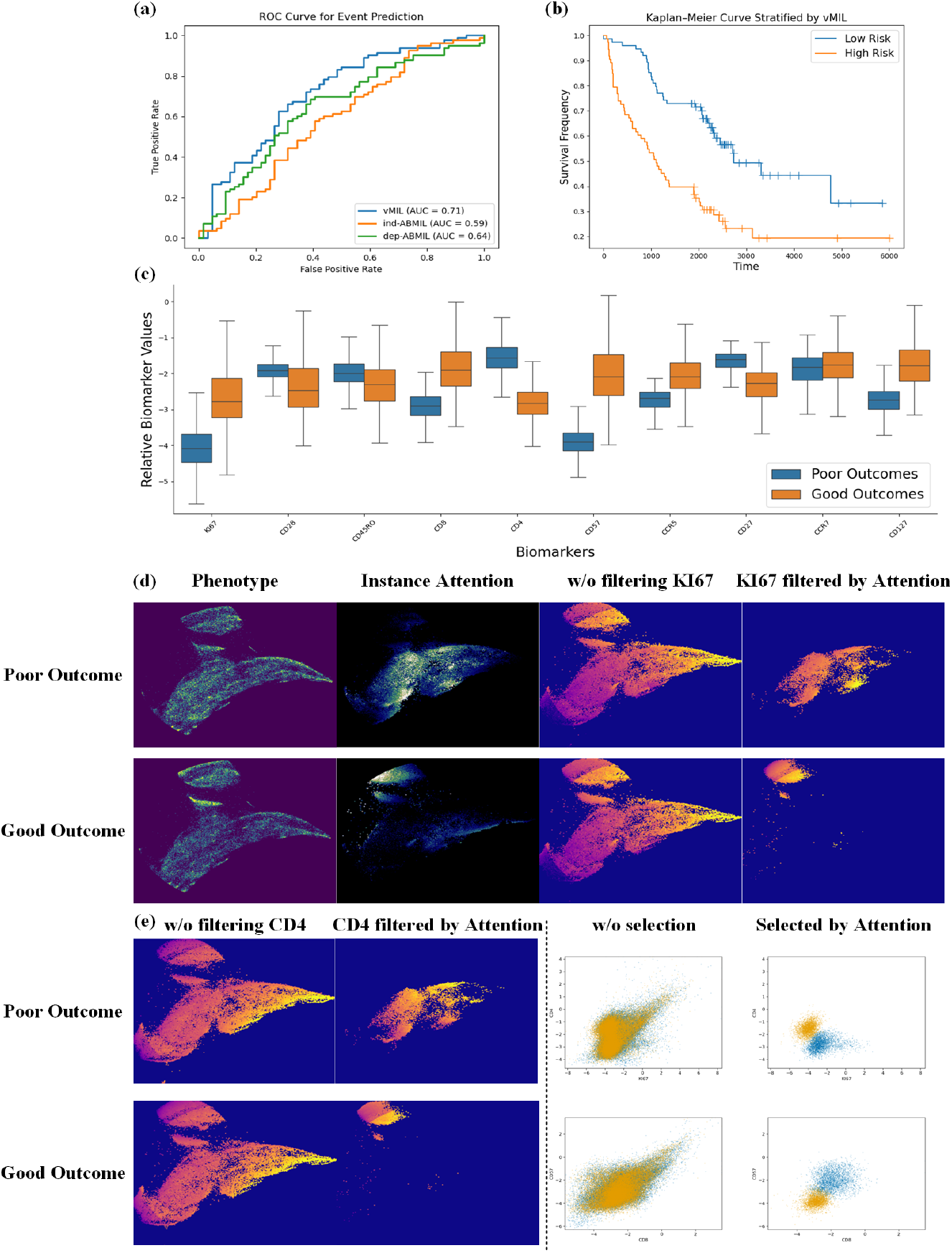
(a) Receiver operating characteristic (ROC) curves for different MIL frame-works. The result differs from the averaged cross-validation result since the curve is plotted based on the collected test set once across splits. (b) Kaplan-Meier curves of patient groups are significantly (log-rank p-value=0.0009) stratified based on vMIL prediction scores. The dataset is stratified evenly into two groups of low and high risks, respectively. (c) Distributions of biomarker reactions for outcome-associated sampled cell populations with high attention scores (top 5%). (d) Visualizations in UMAP spaces. 1^*st*^ column: cell populations sampled from poor and good outcome samples. 2^*nd*^ column: cell populations sensitive to the poor and good outcome attention heads. 3^*rd*^ and 4^*th*^ columns: KI67 response to poor and good outcome associated sampled cell populations without filtering (3^*rd*^) with filtering (top5%, 4^*th*^). (e) Visualizations in UMAP spaces in the first two columns continue for CD4. In the last two columns, cell distributions are presented in selected biomarker space with color representing good outcome (orange) and poor outcome (blue) instance sources. 3^*rd*^ column: Cell population distribution without any selection process in KI67-CD4 and CD8-CD57 spaces. 4^*th*^ column: Cell population distribution selected by network attention (top 5%) in KI67-CD4 and CD8-CD57 spaces.

On the cellular level, MIL enables the selection of phenotype-associated cell populations with distinct biomarker distributions (Wilcoxon test *p* = 0.0019, Fig. 3 (c)) compared to distributions of randomly sampled cells (Supplementary Fig. 8 (a)). However, compared to the TIL dataset, the relationship between phenotypes and cell populations is much more complicated and cannot be identified straightforward in UMAP space (Spearman correlation between two histograms is *ρ*_*s*_ = 0.8347, Fig. 3 (d) 1^*st*^ column). MIL consistently identifies phenotype-associated cell populations across different splits (*ρ*_*s*_ = 0.8394 across splits, see Fig. 3 (d) 2^*nd*^ column for an example and all results in Supplementary Fig. 10). The relationship between captured attention and biomarkers is not straightforward due to the tremendous amount of cells, and direct observations of biomarker reactions without filtering hardly identify meaningful cell populations (Fig. 3 (d) 3^*rd*^ and (e) 1^*st*^ columns, more in Supplementary Fig. 9). The learned MIL attention heads can identify unique cell populations associated with good and poor outcomes (Fig. 3 (d) 4^*th*^ and (e) 2^*nd*^ columns). Moreover, we visualize the cell populations before filtering and after filtering by model attention. Results demonstrate that with model selection, distinctive cell populations are identified and associated with each phenotype (Fig. 3 (e) last two columns). The result suggests that the model identifies good outcome samples through KI67+CD4-CD8+CD57+ cell populations and poor outcome samples through KI67-CD4+CD8-CD57-cell populations.

### MIL uncovers the blast populations in MRD analysis

We utilize the AML dataset [23] to simulate an MRD analysis scenario (Supplementary Fig. 11). Specifically, we follow the established gating process to isolate blasts from eight AML samples, comprising three cytogenetically normal (CN) and five core-binding transcription factor translocation (CBF) samples, alongside healthy bone marrow cells from three samples. These cells are then combined to form a cohort of 256,000 CN blasts, 167,000 CBF blasts, and 387,000 healthy cells. The dataset is partitioned accordingly on the cell level after shuffling to create training, validation, and test sets. However, in simulating an MRD scenario, while the number of blasts is sufficient, the availability of healthy cells is limited. To address this issue, we train a Gaussian Mixture Model (GMM) on the healthy cells in the training set and sample from it to generate additional healthy cells for training samples. This approach enables the creation of an adequately sized MRD dataset with any desired MRD fraction *r*_*d*_.

We focus on vMIL for its intuitive nature in MRD regression circumstances, where assumptions that every cell matters are given. vMIL achieves accurate prediction when *r*_*d*_ is larger than 1 *×* 10^−3^ (averaged mean absolute error (MAE) 5.6 *×* 10^−4^ and averaged mean square error 1.35 *×* 10^−6^, Fig. 4 (a)). If we take a closer look at the situation when *r*_*d*_ ≤ 0.01 (Fig. 4 (b)), the results show that the minimal detectable MRD fraction is 6 *×* 10^−4^, which is the largest predicted *r*_*d*_ when the ground truth is 0. We further check the trained model with a challenge of maximal *r*_*d*_ = 1 *×* 10^−4^ and the result (Supplementary Fig. 14) shows the predicted *r*_*d*_ follows a positive correlation (0.7824) with ground truth.

**Figure 4.**
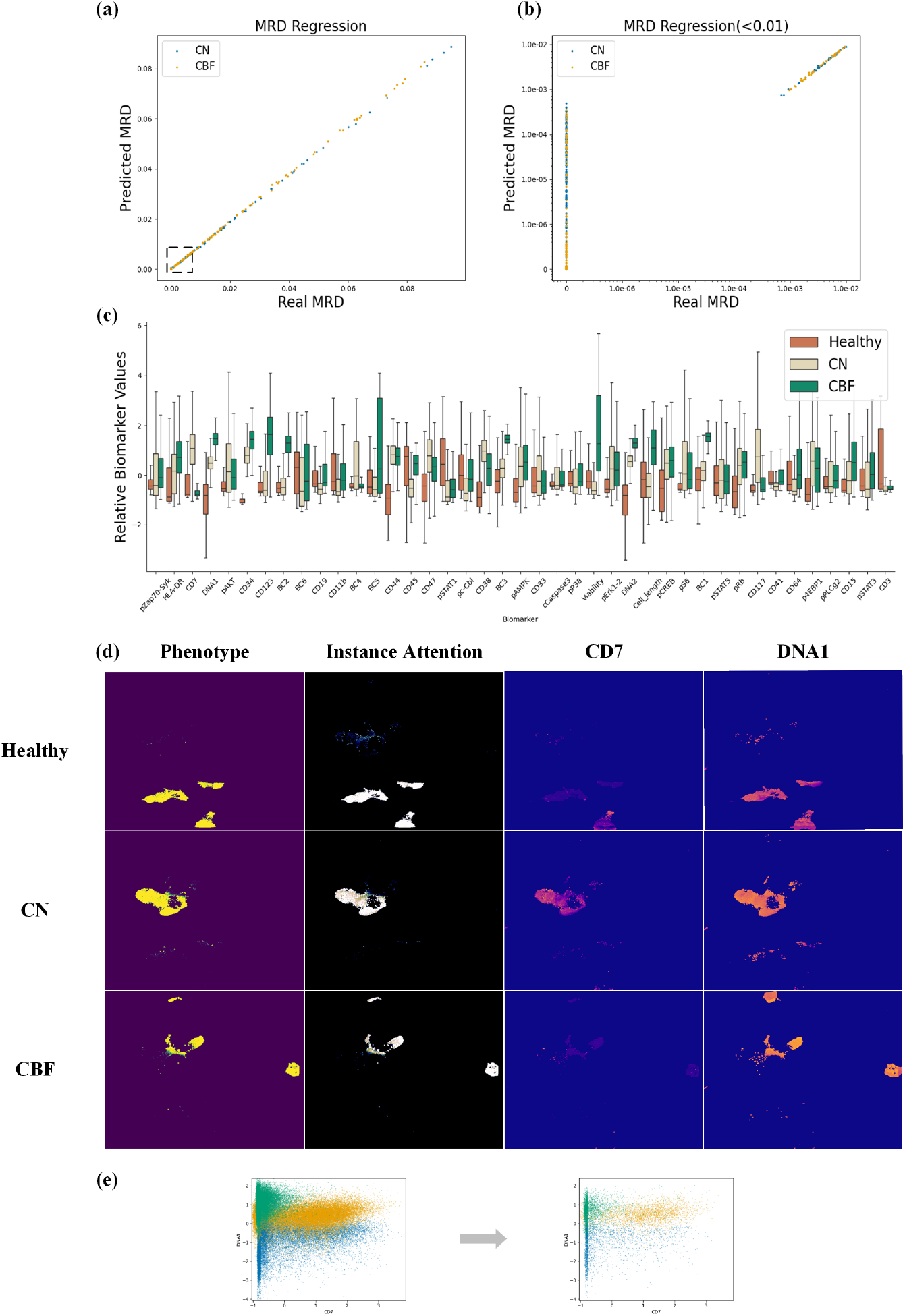
(a,b) Predictive performance for MRD fraction. (c) Distributions of biomarker reactions for cell cohorts with high attention scores (top 5%). (d) UMAP space visualizations for each column are: ground truth, learned instance attention, CD7, and DNA1. (e) Cell population distribution before and after model selection in CD7-DNA1 space.

At the cellular level, MIL enables accurate cell discrimination even though the model itself was never trained with cell-level labels. Using a separate set of cells not included in the simulated experiments (30,000 cells each class), and treating the cellular attention as prediction scores, the model achieved a cellular prediction accuracy of 0.9805, an AUC of 0.9992, and an F1 score of 0.9805. MIL also identifies unmixed subpopulations for MRD prediction. The biomarker reaction of the selected cell population (Fig. 4 (c)) does not show a strong statistical difference from randomly sampled cells (*p* = 0.0475, Supplementary Fig. 13) since a clear biomarker distinction is observable even without filtering. In UMAP space, cell populations with different phenotypes are generally disentangled from each other (Fig. 4 (d) 1^*st*^ column), and this distinctive pattern is perfectly captured by model attention (Fig. 4 (d) 2^*nd*^ column). This disentanglement can be further described by biomarkers CD7 and DNA1 (Fig. 4 (d) 3^*rd*^ and 4^*th*^ columns). Cell populations associated with healthy samples are characterized by CD7-DNA1-, those associated with CN samples by CD7+DNA1_mid_, and those associated with CBF by CD7-DNA1+. Visualization results in CD7-DNA1 space (Fig. 4 (e)) show that the selected subpopulations exhibit less overlap compared to cells without filtering. Compared with vMIL, ABMIL networks’ prediction performance is slightly worse (MAE for ind-ABMIL 1.8 *×* 10^−3^ and for dep-ABMIL 6.1 *×* 10^−3^). Visualization result (Supplementary Fig. 12) shows that ABMILs capture an incomplete subset of blasts.

### MIL identifies COVID severity based on multi-panel FCM data

The COVID dataset [24] comprises four-panel FCM data across varying levels of COVID-19 severity, including 172 healthy, 43 mild, and 54 severe cases. Each panel contains 29 channels, with slight variations in marker composition across panels. The number of cells per panel ranges from 10,000 to 1,000,000.

We first evaluate prediction performance across single-panel and multi-panel settings, focusing on vMIL. Notably, we observe performance variability across panels, with the highest average AUC reaching 0.9573 and the lowest at 0.9295 (Fig. 5 (a)). To integrate information across panels, we design a multi-panel model that averages panel-level features to generate sample-level predictions (details in Architecture details). To mitigate the combinatorial complexity of multi-panel configurations, we introduced panels incrementally, in order of their single-panel performance. Compared to single-panel models, we found that initialization strategies were critical to avoid overfitting. For instance, a randomly initialized multi-panel model achieved a high average AUC of 0.9660 but lower accuracy (0.7915). In contrast, a pretrained multi-panel model initialized with single-panel weights achieves both higher and more stable performance (average AUC of 0.9706 and accuracy of 0.8399; Fig. 5(a)). The best-performing model demonstrates nearly perfect classification for mild and severe cases, although some healthy samples are misclassified as COVID-positive (Fig. 5(b)).

**Figure 5.**
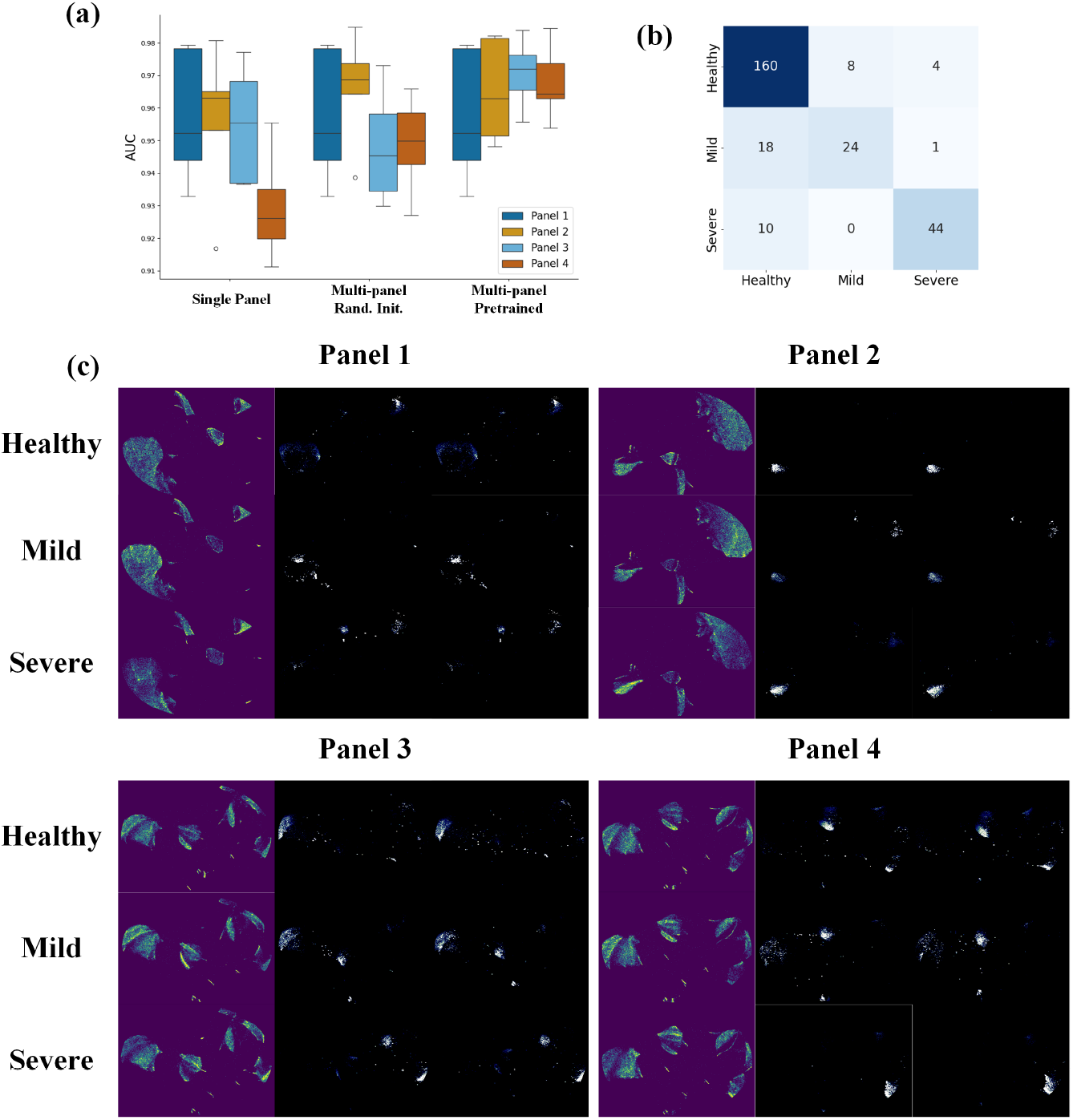
(a) Prediction performance of vMIL on COVID dataset, including models trained on individual panels from random initialization, models trained on multiple panels with random initialization, and models trained on multiple panels initialized with weights from single-panel MIL training. (b) Confusion matrix for the best performance model. (c) UMAP visualizations for each panel, with columns representing cell phenotype, cellular attention from models trained on individual panels, and attention from models trained on multiple panels initialized with single-panel MIL weights.

At the cellular level, we compare the learned representations across panels within the UMAP space (Fig. 5(c)). The phenotype distributions exhibit panel-specific patterns, underscoring the necessity of training models individually for each panel. Within a given panel, phenotypic distinctions are reflected in cell population densities, with a high averaged inter-panel correlation (0.9102). Each panel identifies distinct phenotypes associated with specific immune cell populations. For instance, in panel 2, the healthy phenotype is enriched in CD161+ cells, whereas the severe COVID-19 phenotype is associated with increased frequencies of CD14+ or CD141+ populations (Supplementary Fig. 20). When comparing cell populations identified by models trained on individual panels (Supplementary Fig. 16–19), we observe that the combined panel vMIL model trained from random initialization fails to delineate distinctive cell populations. In contrast, initializing the model with pretrained weights enables the identification of cell populations that closely resemble those discovered by the panel-specific models.

## Discussion

In this study, we apply a series of MIL frameworks across a diverse spectrum of biological and data science contexts, demonstrating their versatility and effectiveness in analyzing high-dimensional flow cytometry data. Compared to the conventional approach of applying separate gating and then using aggregated values for prediction in most cytometry analyses, our MIL framework offers an end-to-end solution for cellular selection based directly on phenotypic information. Specifically, we evaluate our models on clinically relevant tasks, including the characterization of tumor-infiltrating immune cells for cancer subtyping, stratification of HIV patient survival outcomes, prediction of AML minimal residual disease, and assessment of disease severity in COVID-19 patients. Across all settings, our results consistently show that MIL-based approaches not only achieve superior—predictive performance but also provide a level of biological interpretability. By leveraging attention mechanisms within the MIL architecture, our models are able to highlight and quantify the contribution of specific, phenotype-associated cell populations, offering insights into underlying immune mechanisms and potentially actionable biomarkers. This dual capacity for prediction and explanation positions MIL as a powerful tool for both hypothesis generation and clinical decision support in immunological research and precision medicine.

Our results highlight the critical role of network design in both predictive performance and cellular interpretability. The first key component involves the critical settings, including the choice of MIL framework, instance activation strategy, and bag normalization, which directly influence both model accuracy and the cell populations identified. Our observations suggest that using vMIL without bag normalization provides robust performance across most scenarios. While sample-level prediction accuracy is not influenced by the choice of instance activation strategy, this choice alters the distribution patterns of sensitive cell populations, subsequently affecting downstream cellular-level interpretations. The second component involves hyperparameter selection, a critical factor that influences not only the overall predictive performance of the model but also the sensitivity and specificity of the detected cell populations. Using the TIL dataset as a case study, we demonstrate that identifying an optimal network architecture and training configuration is essential for the successful application of MIL frameworks to FCM analysis.

When applying ABMIL, the more commonly used MIL framework in the machine learning community [31–33], the modeling becomes more complicated. For our case, we observe that ABMIL does not necessarily yield improved predictive performance, potentially due to a characteristic of flow cytometry data: the high number of cells per sample relative to the number of measured channels. Furthermore, the attention-based identification of cell populations can probably be countered for the associated class. For example, an attention head trained to focus on class A may inadvertently highlight cell populations relevant to class B, with the sample-level model subsequently calibrating this mismatch. To mitigate this issue, we propose using vMIL-derived pretrained weights to guide attention learning within ABMIL (specifically, ind-ABMIL). We also find that introducing multiple attention heads does not lead to better prediction performance and more interpretable results, which provides further evidence that a more intricate strategy does not necessarily result in better performance. From the COVID dataset, we find that incorporating multiple panels enhances prediction performance by leveraging the complementary information from heterogeneous panels. However, this improvement depends on a well-chosen initialization strategy, such as using pretrained weights from single-panel models, as demonstrated in our case.

In analyzing the identified cell populations, several key findings emerge. MIL frame-works consistently detect stable, phenotype-sensitive cell populations across different training splits, supporting their utility for reliable sample-level prediction. Using the AML dataset, we demonstrate that the MIL frameworks are capable of detecting extremely sparse cell populations with densities as low as 10^−3^, highlighting their sensitivity and effectiveness in identifying rare but potentially clinically significant immune subsets. Notably, vMIL can directly pinpoint phenotype-associated cell subsets, whereas ABMIL models often require additional processing to interpret the results. The patterns of identified cell populations within each dataset are influenced by both the model architecture and the instance activation strategy. Nevertheless, in all cases, the detected populations exhibit clear distinctions between phenotypic groups. These populations can be associated with one or multiple biomarkers, and MIL attention mechanisms offer a powerful, data-driven alternative to traditional gating strategies for identifying phenotype-specific cell subsets. Across datasets, we observe distinct biomarker distribution shifts within the identified sensitive cell populations, except in the AML dataset, where group-level differences emerge even without additional gating beyond preprocessing. Finally, we find that introducing multiple attention heads does not enhance interpretability, suggesting limited benefit for this modification in the context of cell-level analysis.

One limitation of the current framework is that while MIL models can identify phenotype-associated cell populations, they do not capture all such populations. In the TIL dataset, for example, the vMIL model highlights phenotype-distinctive populations associated with CD39, CD127, CD3, and v*α*7.2, whereas the ind-ABMIL model identifies a different set associated with CD103, CD57, CD39, ICOS, and PD-1. In the HIVNH dataset, the presented vMIL model (M1) identifies sensitive cell populations related to KI67, CD4, CD8, and CD57, while another model captures a broader population associated with the entire 10 biomarkers. For the AML dataset, the model detects populations linked to CD7 and DNA1, yet visual inspection of randomly sampled cells reveals additional distinct populations associated with CD34, CD38, and BC2. Similarly, in the COVID dataset, distinct phenotype-specific cell populations emerge in UMAP space for single-panel and combined-panel models. Despite these variations, we reason that the identified cell populations across models are all predictive and biologically meaningful at the sample level. However, the current MIL frameworks tend to focus on specific subsets of informative cells and are not yet capable of capturing the full spectrum of phenotype-relevant cell populations potentially contributing to sample-level classification.

Another limitation of the proposed MIL framework is its inability to handle cases with incomplete channel information. In our experiments, we addressed this by excluding all channels not shared across samples or removing samples that contained unique, non-overlapping FCM channels. This issue leads to a difficult preprocessing process for all datasets. As a result, approximately one-third of the samples in the TIL dataset were excluded based on this criterion. This issue is critical for practical applications, as differences in marker sets across samples, staining protocols, cytometer settings, and patient heterogeneity can all impact the utility of the MIL framework. Therefore, establishing practical benchmarks is strongly encouraged in the future.

The limitation of this study includes comparison with other FCM analytical frame-works in different scenarios, such as cellular clustering strategies [11] and other MRD detection approaches [12].

Leveraging deep learning for FCM analysis presents a promising direction for advancing immune profiling and phenotyping. In contrast to previous approaches, our proposed MIL framework accommodates variable instance sizes across samples and integrates attention mechanisms directly linked to phenotypes, enabling the identification of sensitive cell populations. We systematically benchmark the proposed MIL models across diverse datasets and experimental settings, demonstrating their robustness and effectiveness. We hope that the benchmark datasets and algorithms presented in this work will facilitate further research in this area and contribute to the development and validation of next-generation analytical tools for FCM data.

## Material and Methods

### Dataset details and preprocessing

**TIL dataset** [21] includes mass cytometry data on tumor-infiltrating lymphocytes collected from 144 colorectal or lung cancer patients. For this study, we focus solely on the channels that are complete and exclude those frequently missed by most samples or that lack common channels. After the selection process, we compiled a dataset consisting of 22-channel mass cytometry data from 76 samples, which included 47 cases of colorectal cancer and 29 cases of lung cancer. No additional gating strategies are used for cell selection; instead, we adhere to [21] methodology by applying a logicle transform (t=16409, m=4.5,w=0.25,a=0) followed by a z-score transform to normalize the inputs.

**HIVNH dataset** [22] collected T cell data of 383 HIV-infected individuals. Each sample’s survival time and corresponding censored or uncensored labels are provided per sample. In this study, a classification task is designed to distinguish samples with a survival time of fewer than 4 years and with a survival time of longer than 5 years. Excluding any censored samples, 64 and 83 samples are collected for these two classes, respectively. Following [14], 10 channels except CD3, CD14 and CD19 are used as features. An arcsinh transformation with a cofactor of 150, followed by a z-score transform, is employed to normalize the features.

**AML dataset** for AML bone marrow samples is provided by [23] and we followed the preprecessing process from [14], which includes filtering out AML samples with at least 10% CD34+ blast cells and gathers blasts by gating CD34+CD5- or CD34+CD5mid cell populations. Then the processed samples from both trails are merged into three blast sets based on sample-level classes, which include CN AML (patients SJ10, SJ12, SJ13), CBF [t(8;21) or inv(16)] AML (patients SJ01-SJ05) and healthy controls (H1-H5). Therefore, a dataset of blast cells has been constructed, comprising 256,398 blasts from CN patients, 167,646 CBF blasts, and 387,015 blasts from healthy individuals. All 41 channels are included for the following tasks. To simulate the clinical scenario in which MRD is characterized by extremely small proportions of disease-related blasts, we need to address the challenge of limited blast cells from healthy samples. To overcome this issue (see Supplementary Fig. 11), we first divided our dataset into training, testing, and validation sets based on individual cells. This division is not done on a sample level since the heterogeneity distribution on biomarkers cannot be solved in this small-scale dataset. Subsequently, we trained an additional Gaussian Mixture Model (GMM) with 64 kernels exclusively on healthy blasts from the training set. This model was employed to generate synthetic healthy blasts (Supplementary Fig. 15), thereby augmenting the training set sufficiently for MRD experiments. For validation and testing sets, cells are sampled without replacement in the corresponding cell sets. Following [14], we use the inverse hyperbolic sine transformation with a cofactor of 5. Then, Z-score normalization is applied to each channel to offset the effects of varying magnitudes.

**COVID dataset** [24] comprises flow cytometry data from both healthy individuals and COVID-19 patients, stratified into four clinical severity levels: mild, moderate, severe, and critical. To address class imbalance across these categories, we consolidate the dataset into a three-class classification framework: healthy, mild (a combination of mild and moderate), and severe (a combination of severe and critical). After excluding two samples with incomplete marker panels, the final cohort includes 172 healthy, 43 mild, and 54 severe samples. Each sample is profiled using four distinct panels, with each panel comprising 29 biomarkers that vary across panels. No additional gating or preprocessing steps are applied beyond z-score normalization.

### Architecture details

Three MIL frameworks are introduced in this research for FCM analysis (Fig. 1). At the core of these frameworks is the MLP, composed of stacked linear layers, Softplus activation functions, batch normalization layers, and dropout layers. The output of the MLP is further transformed by an additional activation layer, which modulates the distribution pattern of cell-level attention. In this study, we examine Sigmoid, Softplus, and Softmax activations. For our experiments, we use the dataset 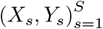, where 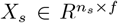 represents the FCM data and *Y*_*s*_ ∈ *R*^*C*^ denotes the sample-level prediction target. The instance size *n*_*s*_ varies across samples, while the number of FCM channels *f* and phenotype classes *C* remain fixed. **vMIL** directly employs an MLP ℳ _1_ to estimate the importance of each cell, taking the processed FCM features as input and outputting corresponding cell-level relevance scores. Specifically, cell-level scores 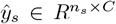 are predicted as *ŷ*_*s*_ = ℳ_1_(*X*_*s*_). Subsequently, sample-level predictions *Ŷ*_*s*_ are obtained by aggregating these cellular scores, i.e., *Ŷ*_*s*_ = 𝒫 (*ŷ*_*s*_). For vMIL, we evaluate two aggregation strategies. The first, referred to as ‘no bag normalization,’ computes the mean of cell-level scores for each of the *C* channels. The second, termed ‘bag normalization,’ sums the cell-level scores within a sample and normalizes the resulting logits across the *C* channels: 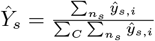. The complete prediction pipeline for vMIL is expressed as *Ŷ*_*s*_ = 𝒫 (ℳ_1_(*X*_*s*_)).

If we interpret the output *ŷ*_*s*_ from *ℳ*_1_ as cellular latent representations, ABMIL introduces an additional MLP *M*_2_ to learn instance-level attention scores for sample-level prediction. This formulation allows 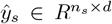 to have a channel dimension *d* that is not constrained to equal *C*. **dep-ABMIL** utilizes the latent variable *ŷ*_*s*_ as input to *ℳ*_2_ to compute instance-level attention scores 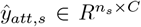, given by *ŷ*_*att,s*_ = ℳ_2_(*ŷ*_*s*_) = ℳ_2_(ℳ_1_(*x*_*s*_)). **ind-ABMIL**, in contrast, uses the raw input *x*_*s*_ as input to ℳ_2_ to generate attention scores directly. Two normalization strategies are applied to the instance attention. The first, ‘no bag normalization,’ normalizes instance attention by dividing by *n*_*s*_. The second, ‘bag normalization,’ normalizes the instance attention across the *C* channels. The resulting normalized attention scores 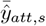 are then used to compute a weighted average of *ŷ*_*s*_, yielding the sample-level feature representation 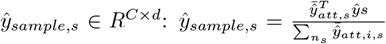. To reduce potential entanglement introduced by a shared sample-level classifier, we employ class-specific MLPs 𝒢 _*c*_ for each of the *C* classes, each producing a one-dimensional prediction. The final sample-level output is constructed by concatenating the class-specific predictions: *Ŷ*_*s*_ = Concat(𝒢 *c*(*ŷ*_*sample,c,s*_)).

For the COVID dataset, **vMIL with multiple panel inputs** is designed to integrate panel-specific predictions by averaging them at the sample level. Rather than relying on aggregated single-panel outputs defined as *Ŷ*_*s,p*_ = 𝒫 (𝒢 _1,*p*_(*X*_*s,p*_)), the final prediction for each sample is computed by averaging across all panels: 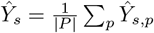. In practice, the multi-panel model structure is flexible and can be adapted by the user as needed.

### Experiment settings

Critical design choices, including the type of MIL framework, instance activation function, and bag normalization strategy, are thoroughly evaluated using the TIL dataset. The insights gained from this analysis are then transferred to guide network design in the remaining three datasets. Hyperparameter tuning is conducted based on experiments across all four datasets (Supplementary Table 2 and 3). The explored hyperparameters include training settings (learning rate and weight decay) as well as network architecture parameters (MLP size and dropout rate). For MLP sizing, we adopt the following strategy: given a network depth *L*, the logarithm of the hidden layer width *W* is sampled between *L* + *V*_*min*_ and *L* + *V*_*max*_, where *V*_*min*_ and *V*_*max*_ are predefined constants greater than 1. Consequently, the width of the *i*-th layer is defined as 2^*L*−*l*+*w*^. For ABMIL, feature extraction and attention networks use independent MLP settings.

Cross-validation is used in all experiments except for the AML dataset. In the remaining three datasets, the dataset is split into train:validation:test = 3:1:1, and fixed 5 cross validation is used. We perform randomized hyperparameter search by sampling multiple combinations from the predefined search space and selecting the model with the lowest validation loss as the optimal configuration. For the TIL dataset, we sample 20 hyperparameter sets for each critical experimental setting. For the HIVNH and COVID datasets (single-panel models), we conduct 10 hyperparameter search trials. For the AML dataset, 5 trials are performed. Specifically, for the COVID dataset, since most biomarkers are shared between panel 1 and panel 2, as well as between panel 3 and panel 4, we perform hyperparameter search only for panel 1 and panel 3. The resulting optimal hyperparameters are then reused for panel 2 and panel 4, respectively.

In the training process, we use the ADAM optimizer (*β*_1_ = 0.9 and *β*_2_ = 0.999) with clipnorm 1 and binary cross entropy as the training target. Early-stopping criterion is introduced to stop training if the validation loss does not improve for specific epochs (Supplementary Table 7). For the HIVNH and COVID datasets, an instance sampling strategy is employed to address the large number of instances per sample. We verified that this sampling approach does not compromise prediction performance.

### Statistical analysis

To assess group-level differences, we use the t-test for comparisons involving two groups and one-way ANOVA for multiple-group comparisons, with assumptions of independence of observations, approximate normality of residuals within each group, and homogeneity of variances across groups. The log-rank test for Kaplan–Meier survival curves assumes independent groups, non-informative censoring, and approximately proportional hazards over time. The Wilcoxon rank-sum test evaluates group-level differences under the assumptions of independent samples and similarly shaped distributions across groups. Spearman correlation is employed to measure histogram similarity.

### Visualization settings

UMAP visualizations are constructed following the approach described in [34]. For each attention head, a normalized histogram is generated by dividing the attention-weighted histogram in UMAP space by the corresponding unweighted histogram. To enhance interpretability, the resulting figures are smoothed using a Gaussian filter with a standard deviation of 1. For biomarker activity visualization in UMAP space, values are normalized by subtracting the minimum value of each respective biomarker. Gating plots are generated as scatter plots, with biomarker intensities scaled using min-max normalization and further clipped at the 0.75 quantile to reduce the impact of outliers.

## Data Availability

All datasets used in this study are publicly available through FlowRepository or Cytobank databases. The TIL dataset^1^(FR-FCM-ZYWM), HIVNH dataset^2^(FR-FCM-ZZZK), and COVID dataset^3^(FR-FCM-Z5PC) can be accessed via FlowRepository. The AML dataset^4^ is available through Cytobank.

## Code Availability

The experiment framework, named FlowMIL, can be found at https://github.com/Zhiyuan-Ding/FlowMIL

## Author Contributions

A.B. and Z.D. conceptualize the framework. Z.D. conducts the experiments and writes the manuscript. A.B. reviews the results and the manuscript.

## Footnotes

1 https://flowrepository.org/id/FR-FCM-ZYWM

2 https://flowrepository.org/id/FR-FCM-ZZZK

3 https://flowrepository.org/id/FR-FCM-Z5PC

4 https://community.cytobank.org/cytobank/experiments/44185

