## Supplementary materials for "Application and Characterization of the Multiple Instance Learning Framework in Flow Cytometry"

### Supplementary Information for

### Understanding Multi-Instance Learning for Flow Cytometry Analysis

#### Supplementary Notes

**Introduced tricks in MIL FCM analysis.** In this part, we introduce the special network designs we made for FCM analysis.

**Phenotype associated attention heads.** To ensure that the selected cell populations are directly associated with specific phenotypes, each attention head in both vMIL and ABMIL is explicitly aligned with one of the sample-level classes.

**Isolating the sample-level network structure.** To minimize the influence of the sample-level network on instance-level interpretation, we design the sample-level components to be as lightweight as possible. Additionally, the learnable networks corresponding to each phenotype are made as disentangled as possible, reducing interference between class-specific features. We find that this architectural choice significantly impacts instance-level interpretability. Without such separation, the model tends to bypass the instance-level MLP, instead learning to make predictions by directly comparing aggregated sample-level features, resulting in poor or non-informative cell-level insights. In the case of vMIL, we avoid using any bag-level MLPs entirely, relying instead on the direct aggregation of sample-level logits for prediction. For ABMIL, class-specific features are each processed through their own sample-level MLP, and are only combined at the final output layer, maintaining separation throughout most of the network.

**Using vMIL to initialize dep-ABMIL.** Even with deliberate architectural design to minimize the influence of the sample-level network on instance-level interpretation, we observe that ABMIL can still learn countered instance attention. In such cases, the model predicts a given phenotype by focusing on cell populations associated with other phenotypes rather than those directly relevant, compromising interpretability. This phenomenon does not occur in the vMIL due to the direct correspondence between attention logits and sample-level predictions, which enforces a more faithful alignment between phenotype-specific features and cell populations. Motivated by this observation, we propose using the trained weights from vMIL as initialization for ABMIL models, leading to the design of dep-ABMIL, a variant that can directly load and fine-tune from vMIL weights to enhance phenotype-specific attention alignment and interpretability.

**Instance sampling in training process.** To enhance training robustness and efficiency, we introduce an instance sampling strategy that randomly selects a subset of instances from each sample during training, rather than using the full set of cells. Although this approach does not lead to a measurable improvement in predictive performance, it also does not result in any degradation. Importantly, it enables scalable training on extremely large datasets, such as the HIVNH and COVID cohorts, while significantly reducing computational load and training time.

**Introducing extra channels for Softmax Activation.** Softmax enforces that the attention weights assigned to instances across phenotypes sum to 1, which can be overly restrictive for FCM analysis, where cells may not correspond clearly to any specific phenotype. To address this limitation, and inspired by [1], we introduce an additional attention channel that does not correspond to any phenotype. This extra channel allows the model to account for non-informative or ambiguous cells, thereby enabling the learning of sparse, phenotype-sensitive cell populations while still using the Softmax activation function.

**Instance activations distribution assumption.** Applying different instance activation functions reflects distinct underlying assumptions about the resulting probability distribution (PD) over instances. In this discussion, we focus on the three activation functions used in our framework: Sigmoid, Softplus, and Softmax. A central distinction among these functions lies in whether the PD is considered proper, that is, whether the attention weights assigned to all instances (cells) sum to 1. Softmax enforces a proper PD, ensuring that the total attention is normalized across instances. In contrast, Sigmoid and Softplus produce improper PDs, allowing for cells to receive attention values independently. This permits some cells to be associated with multiple phenotypes or none at all—better reflecting complex biological systems where such

| Model | AUC | ACC | F1 |
|---|---|---|---|
| PACMAN | 0.9452 | 0.8877 | 0.8950 |
| FlowSOM | 0.8695 | 0.7106 | 0.7516 |
| FlowClust | 0.8891 | 0.7571 | 0.7891 |
| Depeche | 0.9261 | 0.9039 | 0.9091 |
| MIL-V0 | 0.9545 | 0.9179 | 0.9208 |
| dep-ABMIL | 0.9524 | **0.9212** | **0.9225** |
| ind-ABMIL | **0.9602** | 0.8899 | 0.8950 |

Table 1: Comparison of cell clustering strategies on TIL dataset

overlaps naturally occur. While Sigmoid and Softplus are theoretically capable of producing equivalent attention patterns, they differ in the shapes of their activation curves, which influence the resulting probability density. In practice, due to stochastic optimization techniques and model training dynamics, the attention distributions learned using Sigmoid and Softplus may diverge, leading to distinct histogram patterns and potentially different biological interpretations.

**Related Work.** Deep learning frameworks have been applied in FCM analysis. Early-stage works [2, 3] focus on using Convolutional Neural Networks (CNNs) for FCM analysis, where 1D convolutional layers are used to merge biomarker features. For cellular-level interpretation, [3] uses filter-averaged activations for model explanation, while [2] introduces a decision-tree approach to interpret the detected sensitive cell populations. In our work, the stack of convolutional layers is replaced by instance-level MLPs, which directly combine all biomarker features without relying on spatial convolution. One recent work [4] applies ABMIL with two panels to characterize AML-related tasks, establishing a two-stage framework consisting of single-panel pretraining followed by supervised training. We evaluate both the pretraining strategy and the specific sinusoidal encoding proposed in this work, but find that these techniques do not yield performance improvements on the TIL dataset. We attribute this to the relatively smaller sample size of our dataset compared to the cohort used in [4], which reflects a common limitation in real-world FCM applications. Biomarker importance in their work is assessed using predictive power scores, a statistical measure comparing biomarker values between model-selected cells and randomly sampled cells. This is conceptually similar to the biomarker reactivity plots we developed for the TIL, HIVNH, and AML datasets. Other works [5, 6, 7] explore alternative network architectures for FCM analysis, such as set transformers [8] and graph neural networks. However, these approaches often rely on overly complex architectures or require fixed-size sampling of instances, which we consider out of scope for the current study and leave for future work.

### Supplementary Tables

| Hyperparameter | TIL | HIVNH | AML | COVID |
|---|---|---|---|---|
| MLP Block Size | $(2, 4)$ | $(2, 3)$ | $(2, 3)$ | $(2, 4)$ |
| Max-Width | $2^{(L+1,L+4)}$ | $2^{(L+1,L+4)}$ | $2^{(L+1,L+4)}$ | $2^{(L+1,L+3)}$ |
| Dropout | $(0, 0.2)$ | $(0, 0.2)$ | $(0, 0.2)$ | $(0, 0.4)$ |
| Learning Rate | $(5 \times 10^{-4}, 10^{-2})$ | $(10^{-3}, 2 \times 10^{-2})$ | $(10^{-3}, 2 \times 10^{-2})$ | $(5 \times 10^{-4}, 10^{-2})$ |
| Weight Decay | $(0, 10^{-3})$ | $(0, 10^{-3})$ | $(0, 10^{-3})$ | $(0, 10^{-3})$ |

Table 2: Hyperparameter searching spaces for vMIL.

| Hyperparameter | TIL | HIVNH | AML |
|---|---|---|---|
| F.E. Block Size | $(2, 4)$ | $(2, 3)$ | $(2, 4)$ |
| F.E. Max-Width | $2^{(L+1, L+4)}$ | $2^{(L+1, L+3)}$ | $2^{(L+1, L+3)}$ |
| Att. Block Size | $(1, 3)$ | $(1, 3)$ | $(1, 3)$ |
| Att. Max-Width | $2^{(L+1, L+4)}$ | $2^{(L+1, L+4)}$ | $2^{(L+1, L+3)}$ |
| Dropout | $(0, 0.2)$ | $(0, 0.2)$ | $(0, 0.4)$ |
| Learning Rate | $(5 \times 10^{-4}, 10^{-2})$ | $(10^{-3}, 2 \times 10^{-2})$ | $(5 \times 10^{-4}, 10^{-2})$ |
| Weight Decay | $(0, 10^{-3})$ | $(0, 10^{-3})$ | $(0, 10^{-3})$ |

Table 3: Hyperparameter searching spaces for ABMIL. Hyperparameter searching space is the same for dep-ABMIL and ind-ABMIL. F.E. represents feature extraction MLP. Att. represents attention MLP. COVID dataset is not included since ABMIL is not used in experiment.

| Dataset | Inst. Act. | Bag Norm. | MLP Block Size | Max-Width | Dropout | Learning Rate | Weight Decay |
|---|---|---|---|---|---|---|---|
| TIL | SG | - | 4 | 128 | 0 | 0.005 | $10^{-5}$ |
| TIL | SP | - | 4 | 32 | 0 | 0.01 | $10^{-4}$ |
| TIL | SM | - | 4 | 128 | 0.1 | 0.005 | $10^{-5}$ |
| TIL | SG | + | 2 | 32 | 0.2 | 0.001 | $10^{-5}$ |
| TIL | SP | + | 2 | 16 | 0.1 | 0.001 | $10^{-3}$ |
| TIL | SM | + | 2 | 32 | 0.1 | 0.001 | $10^{-3}$ |
| HIVNH | SG | + | 2 | 32 | 0.2 | 0.02 | 0 |
| AML | SG | + | 3 | 16 | 0.2 | 0.002 | $10^{-4}$ |
| COVID-1 | SG | + | 3 | 16 | 0.1 | 0.02 | 0 |
| COVID-2 | SG | + | 2 | 16 | 0 | 0.002 | $10^{-5}$ |

Table 4: Settings for vMIL with best performances. Inst. Act. represents instance activation. Bag Norm. represents the bag normalization strategy. SG, SP, and SM represent Sigmoid, Softplus and Softmax. COVID-1 represents the searched setting for COVID dataset panel 1 (BDC-CR1) and panel 2 (BDC-CR2). COVID-2 represents the searched setting for COVID dataset panel 3 (TNK-CR1) and panel 4 (TNK-CR2).

| Dataset | Inst. Act. | Bag Norm. | F.E. Block Size | F.E. Max-Width | Att. Block Size | Att. Max-Width | Dropout | Learning Rate | Weight Decay |
|---|---|---|---|---|---|---|---|---|---|
| TIL | SG | - | 2 | 32 | 2 | 32 | 0.1 | 0.0005 | $10^{-4}$ |
| TIL | SP | - | 4 | 32 | 1 | 16 | 0 | 0.001 | 0 |
| TIL | SM | - | 4 | 128 | 3 | 16 | 0.2 | 0.0005 | $10^{-5}$ |
| TIL | SG | + | 3 | 64 | 1 | 8 | 0.2 | 0.001 | $10^{-3}$ |
| TIL | SP | + | 4 | 128 | 1 | 8 | 0 | 0.005 | 0 |
| TIL | SM | + | 3 | 64 | 3 | 16 | 0.1 | 0.005 | $10^{-4}$ |
| HIVNH | SG | - | 2 | 32 | 2 | 8 | 0.2 | 0.005 | $10^{-4}$ |
| AML | SG | + | 2 | 8 | 3 | 16 | 0.3 | 0.002 | $10^{-4}$ |

Table 5: Settings for dep-ABMIL with best performances. Inst. Act. represents instance activation of the attention network. Bag Norm. represents the bag normalization strategy. F.E represent feature extraction MLP. Att. represents attention MLP. SG, SP, and SM represent Sigmoid, Softplus and Softmax.

| Dataset | Inst. Act. | Bag Norm. | F.E. Block Size | F.E. Max-Width | Att. Block Size | Att. Max-Width | Dropout | Learning Rate | Weight Decay |
|---|---|---|---|---|---|---|---|---|---|
| TIL | SG | - | 2 | 32 | 3 | 32 | 0 | 0.005 | $10^{-5}$ |
| TIL | SP | - | 2 | 16 | 2 | 16 | 0.1 | 0.005 | $10^{-4}$ |
| TIL | SM | - | 3 | 64 | 3 | 32 | 0.1 | 0.0005 | 0 |
| TIL | SG | + | 2 | 32 | 3 | 16 | 0 | 0.005 | $10^{-4}$ |
| TIL | SP | + | 2 | 16 | 2 | 8 | 0.1 | 0.005 | $10^{-3}$ |
| TIL | SM | + | 2 | 32 | 3 | 16 | 0.1 | 0.002 | $10^{-3}$ |
| HIVNH | SG | - | 2 | 32 | 3 | 32 | 0.1 | 0.005 | $10^{-5}$ |
| AML | SG | + | 1 | 8 | 2 | 16 | 0.3 | 0.005 | $10^{-3}$ |

Table 6: Settings for ind-ABMIL with best performances. Inst. Act. represents instance activation of the attention network. Bag Norm. represents the bag normalization strategy. F.E represent feature extraction MLP. Att. represents attention MLP. SG, SP, and SM represent Sigmoid, Softplus and Softmax.

| Hyperparameter | TIL | HIVNH | AML | COVID-1 | COVID-2 | COVID-3 |
|---|---|---|---|---|---|---|
| Patience | 300 | 50 | 50 | 50 | 50 | 5 |
| Maximal Epochs | 3000 | 200 | 800 | 200 | 200 | 50 |
| Batch Size | 32 | 32 | 32 | 32 | 32 | 32 |
| Inst. Samp. | - | + | - | - | + | + |

Table 7: Training profiles for MIL frameworks across datasets. COVID-1 represents the single panel model. COVID-2 represents combined-panel model with random initialization. COVID-3 represents combined-panel model initialized with single-panel weights. Inst. Samp. represents instance sampling.

### Supplementary Figures

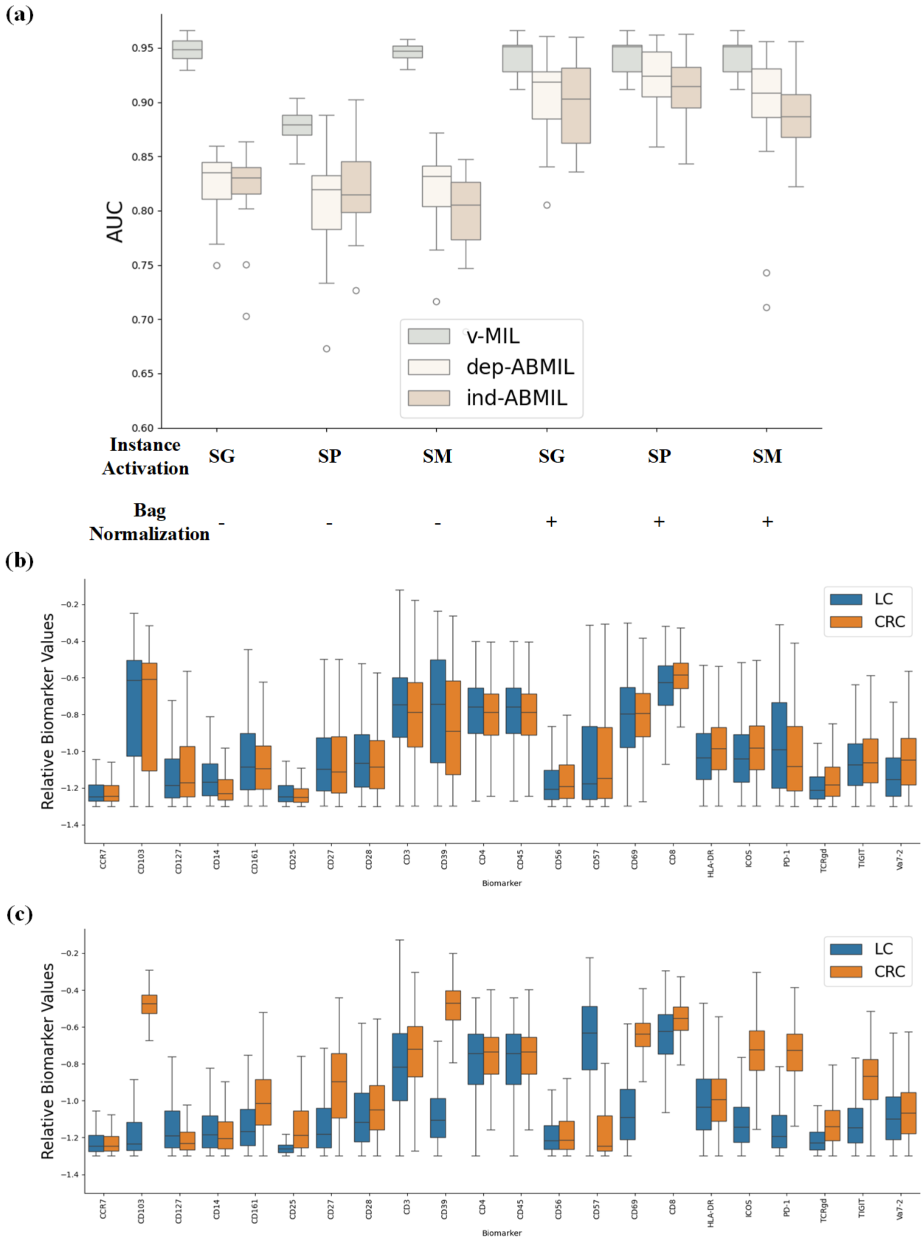


Figure 1: (a) Averaged CRC and LC subtyping performance across different MIL framework settings, including MIL structures, instance activation, and bag normalization strategies with different hyperparameter settings, including training profiles and network hyperparameters. The evaluated instance activation methods include Sigmoid (SG), Softplus (SP), and Softmax (SM). Bag normalization methods across attention heads are combined with specific aggregation strategies: averaging across cells without bag normalization (−) and summing across cells with bag normalization (+). (b) Biomarker values are statistically analyzed for cell populations with random sampling, derived from CRC and LC sampled cells. (c) Biomarker values are statistically analyzed for sensitive cell populations of ind-ABMIL with Sigmoid activation and bag normalization.

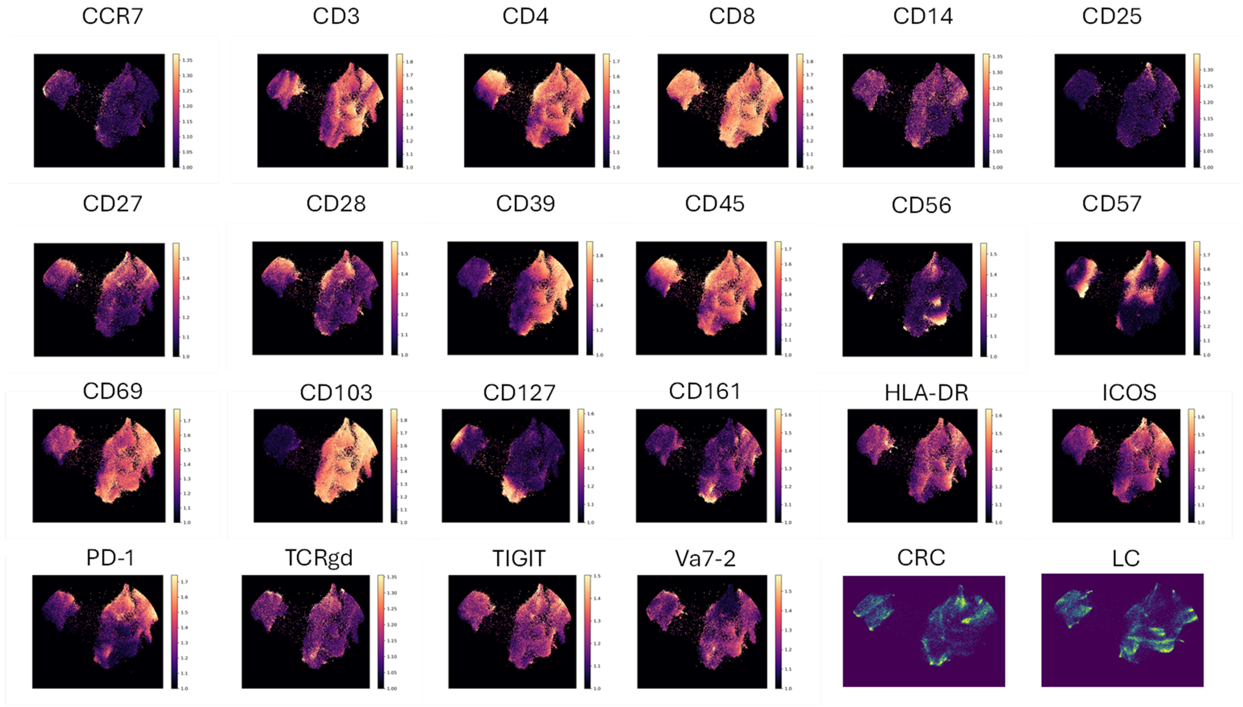


Figure 2: Biomarker activations for all of the 22 channels of TIL datasets in UMAP space with 30,000 cells sampled from CRC and LC samples respectively.

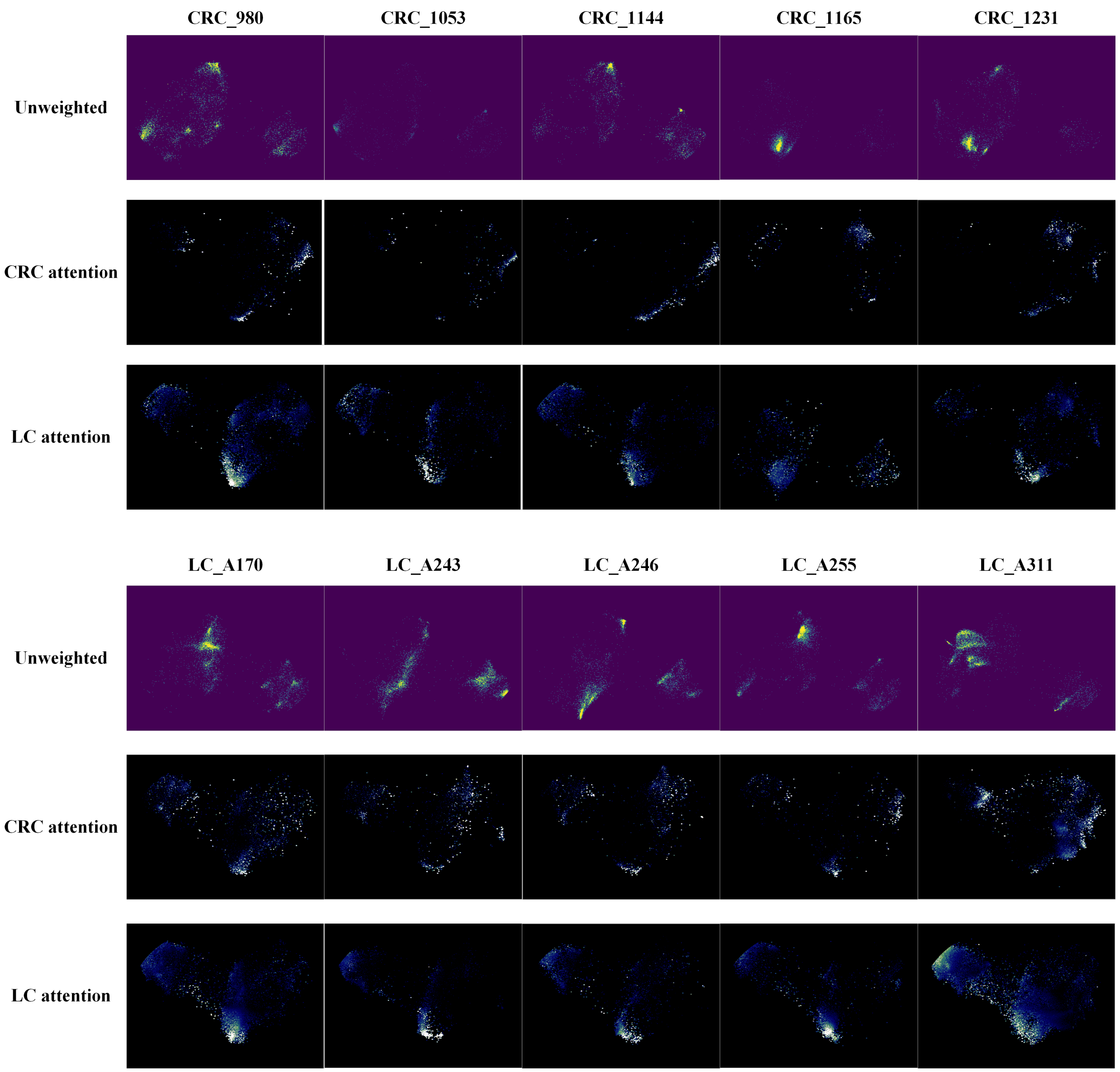


Figure 3: $1^{st}$ and $4^{th}$ rows: Selected sample cell population distribution in UMAP space.5 LC and CRC samples are selected randomly from the TIL dataset with more than 5,000 cells. $2^{nd}$ and $5^{th}$ rows: cell populations with high CRC-associated attention from model M1. $3^{rd}$ and $6^{th}$ rows: cell populations with high LC-associated attention from model M1.

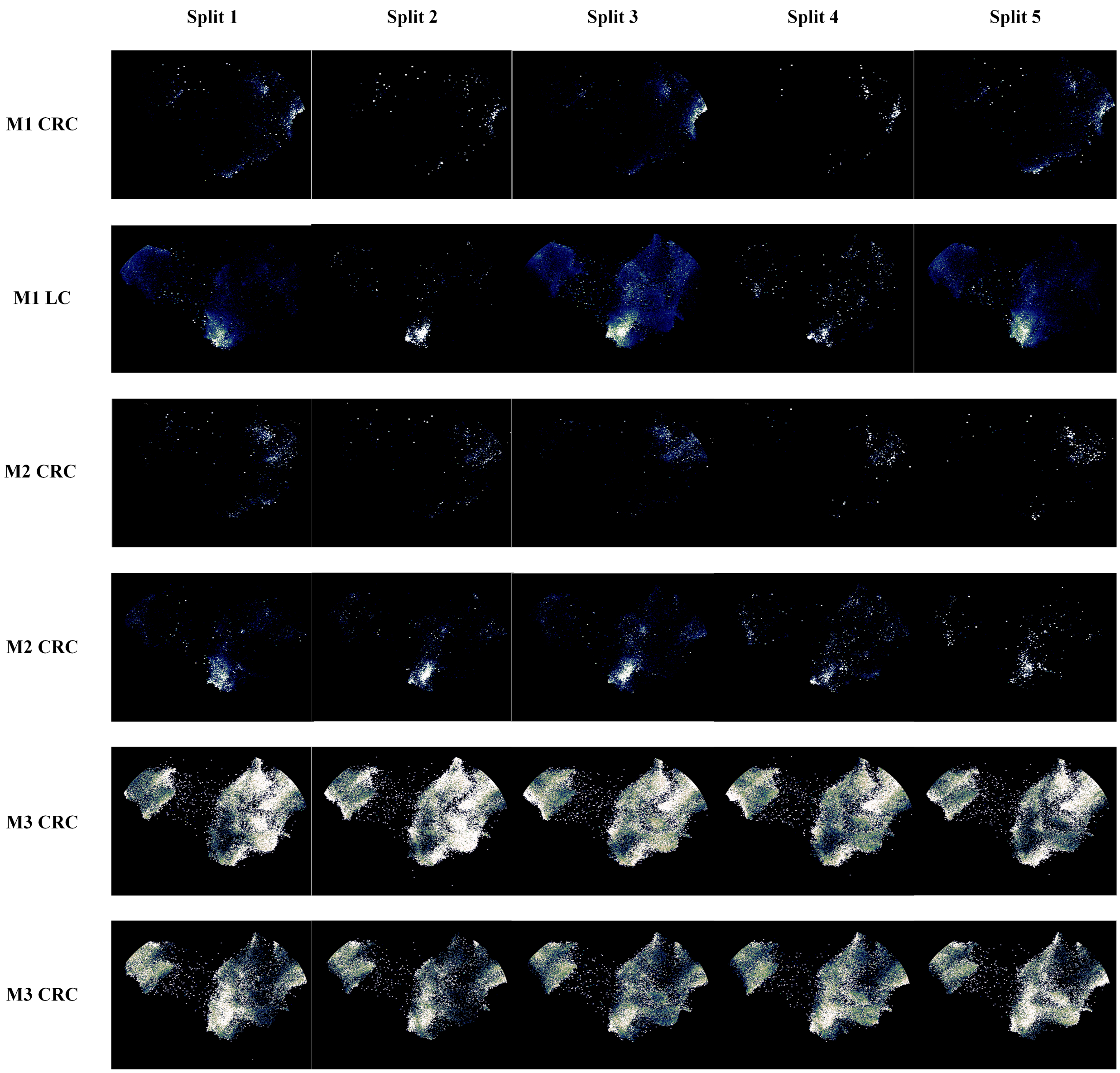


Figure 4: Cell population with high CRC and LC associated attention values from models. M1: vMIL with Sigmoid instance activation and bag normalization. M2: vMIL with Softplus instance activation and bag normalization. M3: vMIL with Sigmoid instance activation and without bag normalization. Observations: Cell population distribution patterns across different experiment splits are similar. M1 and M2 share similar cell population distribution patterns while M3 does not with either M1 or M2.

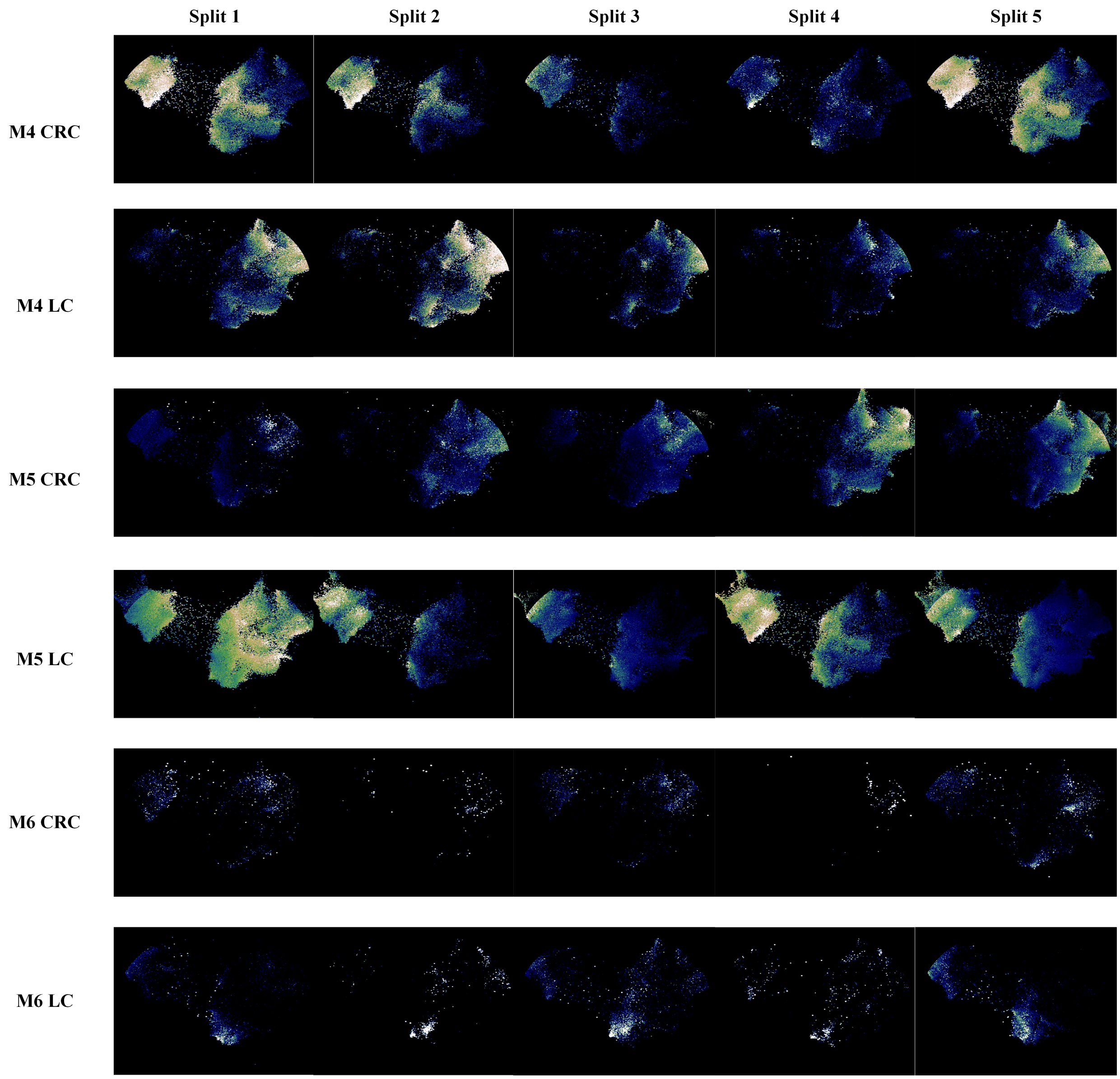


Figure 5: Cell populations with high CRC- and LC-associated attention values were analyzed across different models. M4 corresponds to ind-ABMIL with Sigmoid instance activation and no bag normalization. M5 represents dep-ABMIL with Softplus instance activation and bag normalization. M6 is ind-ABMIL with Sigmoid instance activation and no bag normalization but initialized with vMIL weights. Observations show that M4 and M5 exhibit 'opposite attention' distributions. M4 follows a pattern similar to M3 and includes the same sensitive cell populations as M1 and M2, making it the positive model. In contrast, M5 is considered the negative model. In M6, using pretrained weights from the corresponding vMIL leads to consistent attention distributions and mitigates the 'opposite attention' issue.

(a)

(b)

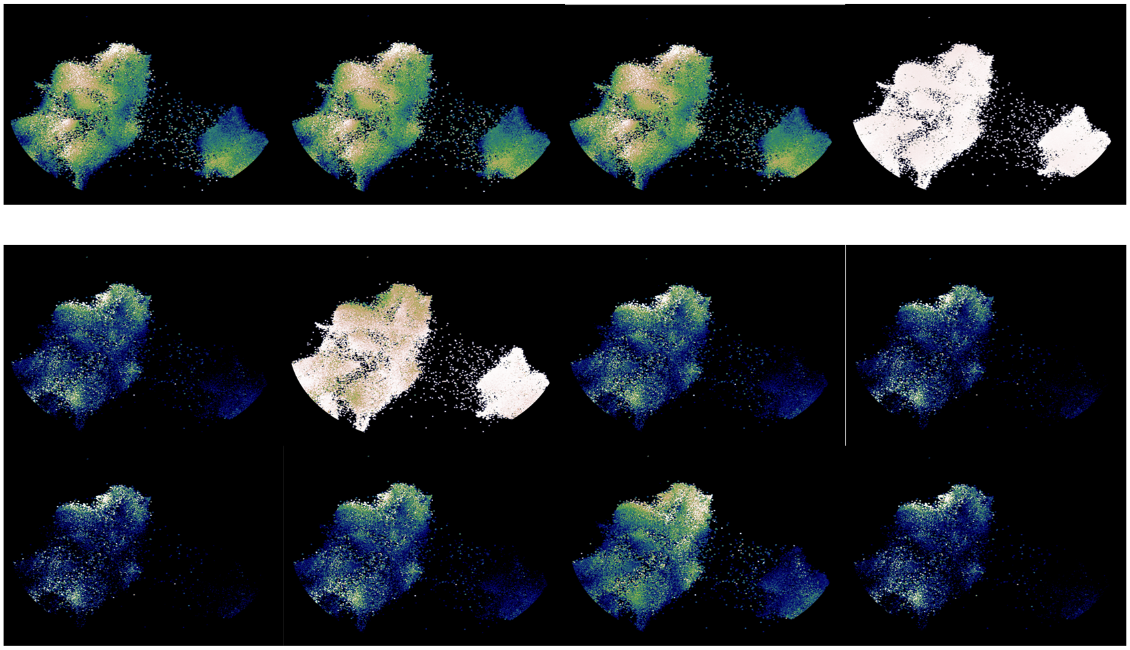

Figure 6: Sensitive cell populations identified by ABMIL-MH from different attention heads for (a) 4 heads and (b) 8 heads.

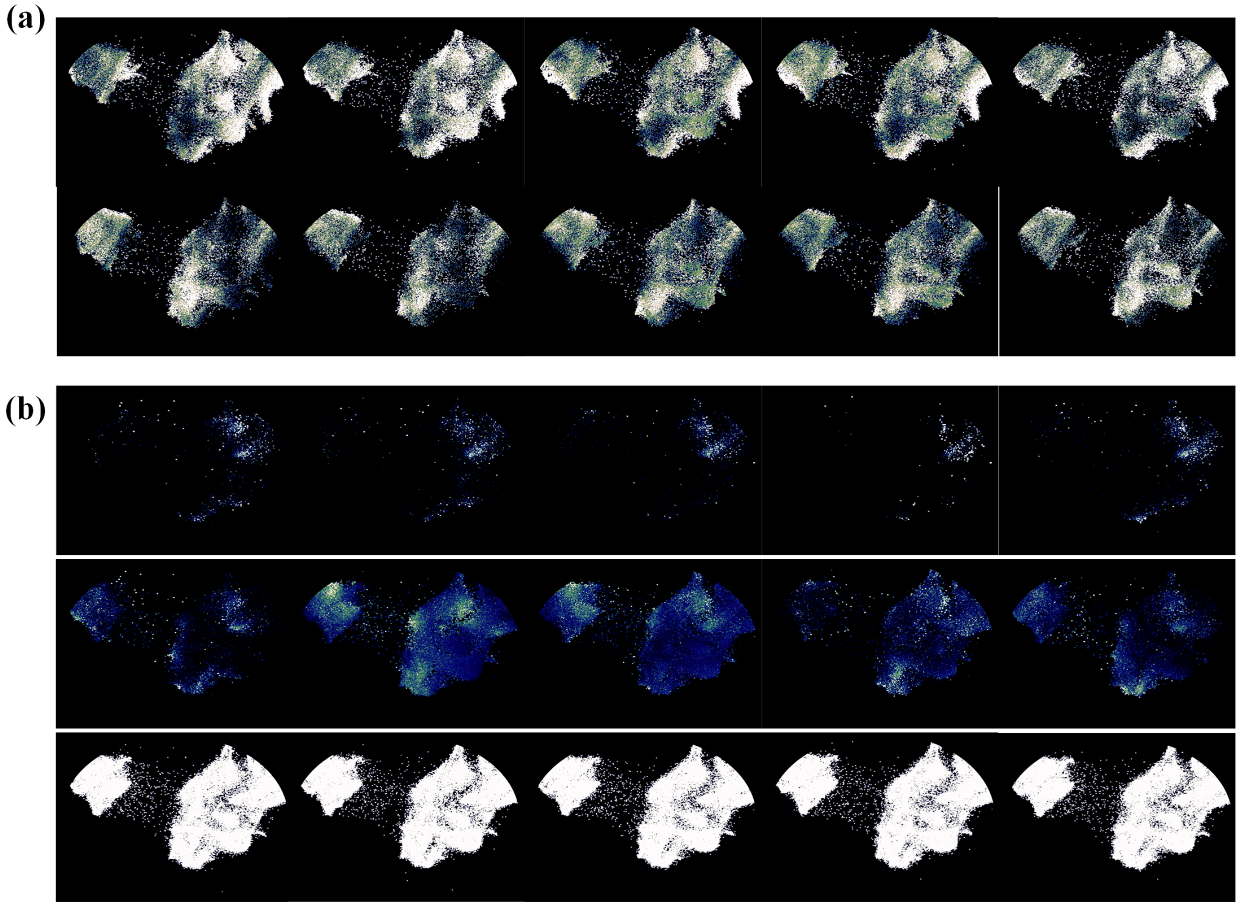


Figure 7: Sensitive cell populations in UMAP space for vMIL with Softmax activation. (a) The number of attention heads is set to 2, corresponding to the number of phenotypes. (b) An additional, unassigned attention head is introduced (third row), which is not associated with any specific phenotype. This strategy enables the model to identify sparse, phenotype-specific cell populations more effectively.

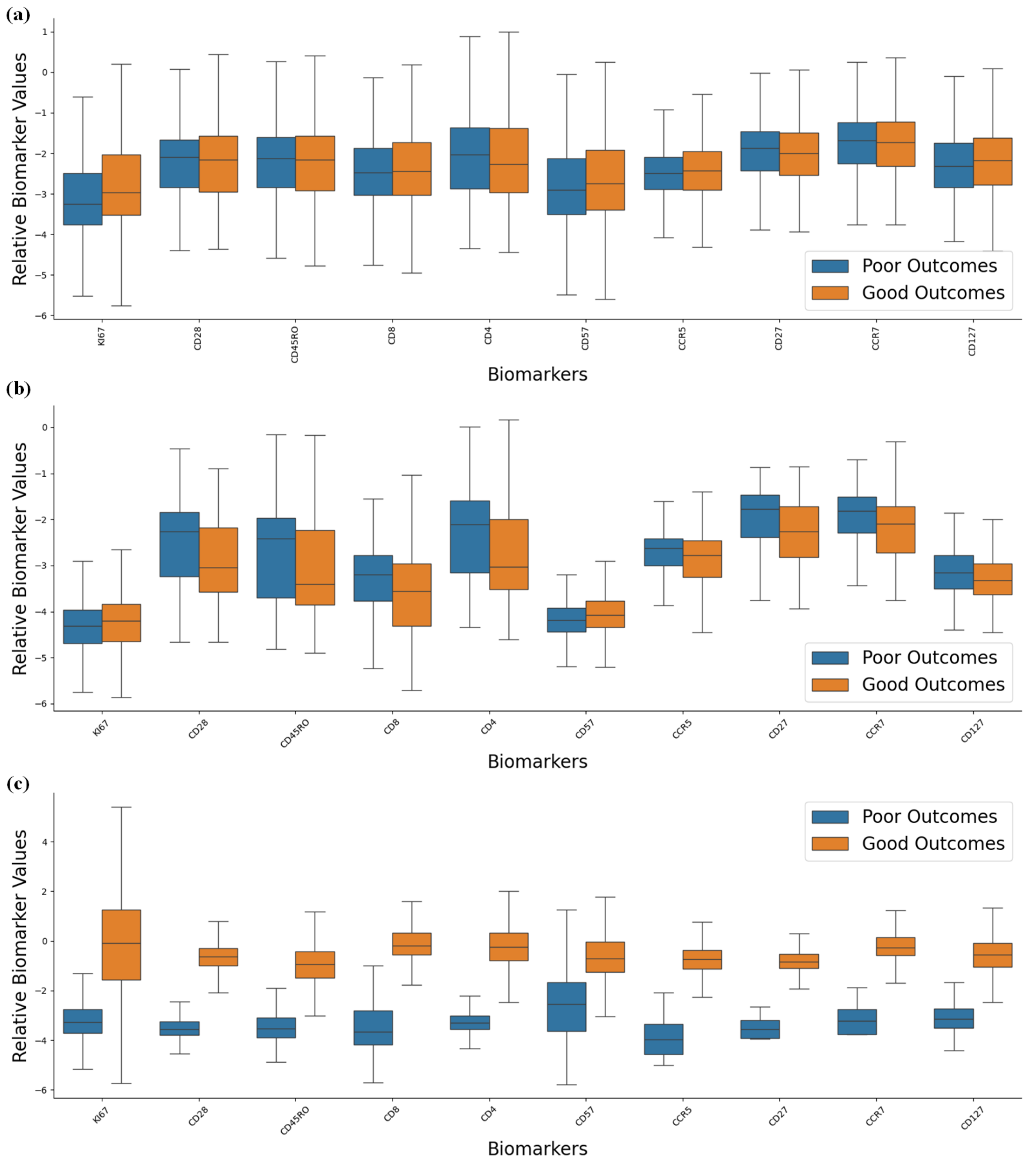


Figure 8: Biomarker values are statistically analyzed for cell populations with (a) random sampling, (b) vMIL M1, and (c) vMIL M2, derived from good and poor outcomes associated sampled cells.

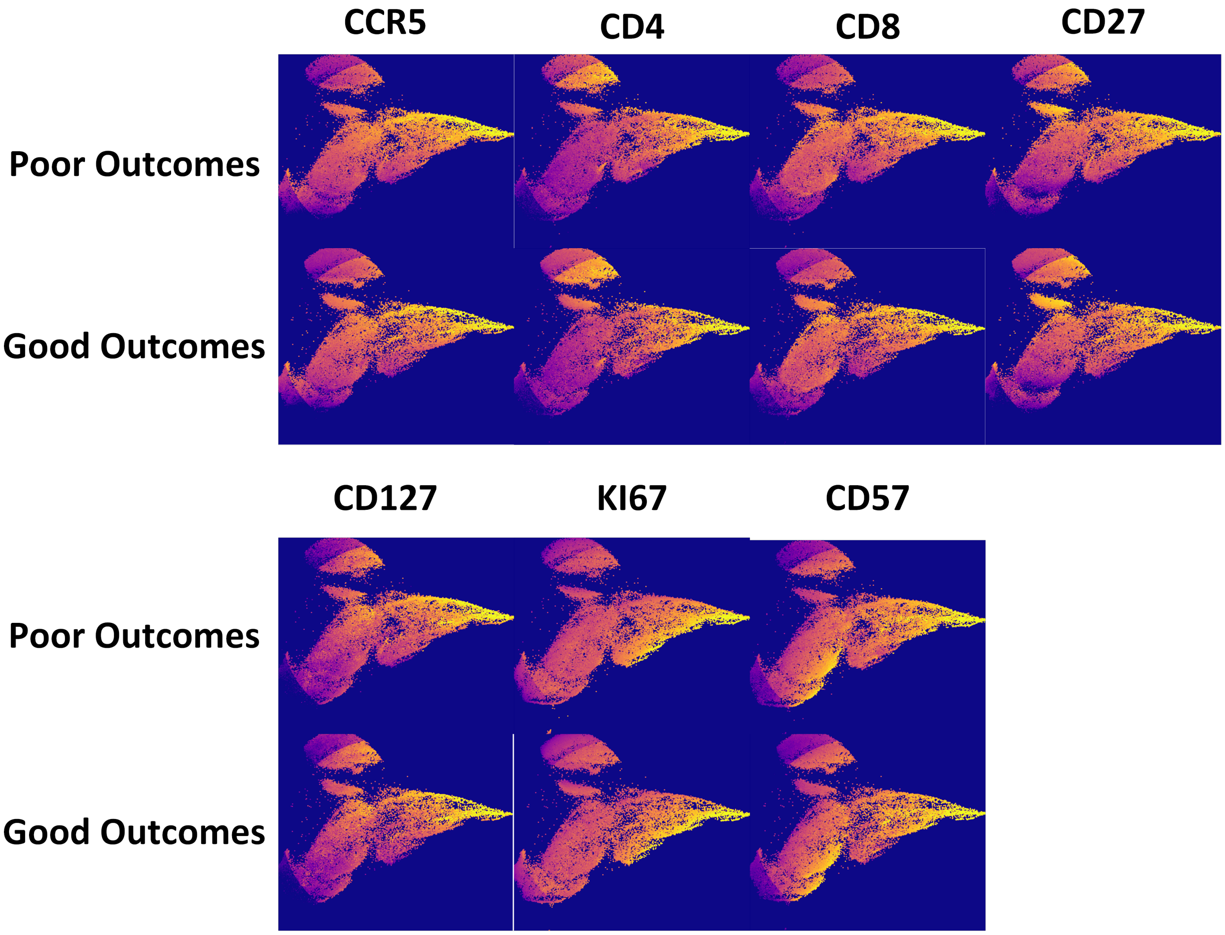


Figure 9: Biomarker reactions are visualized in UMAP space for HIVNH cell populations sampled from good and poor outcomes associated sampled cells. Biomarkers are selected based on groupwise differences in high-attention cell populations.

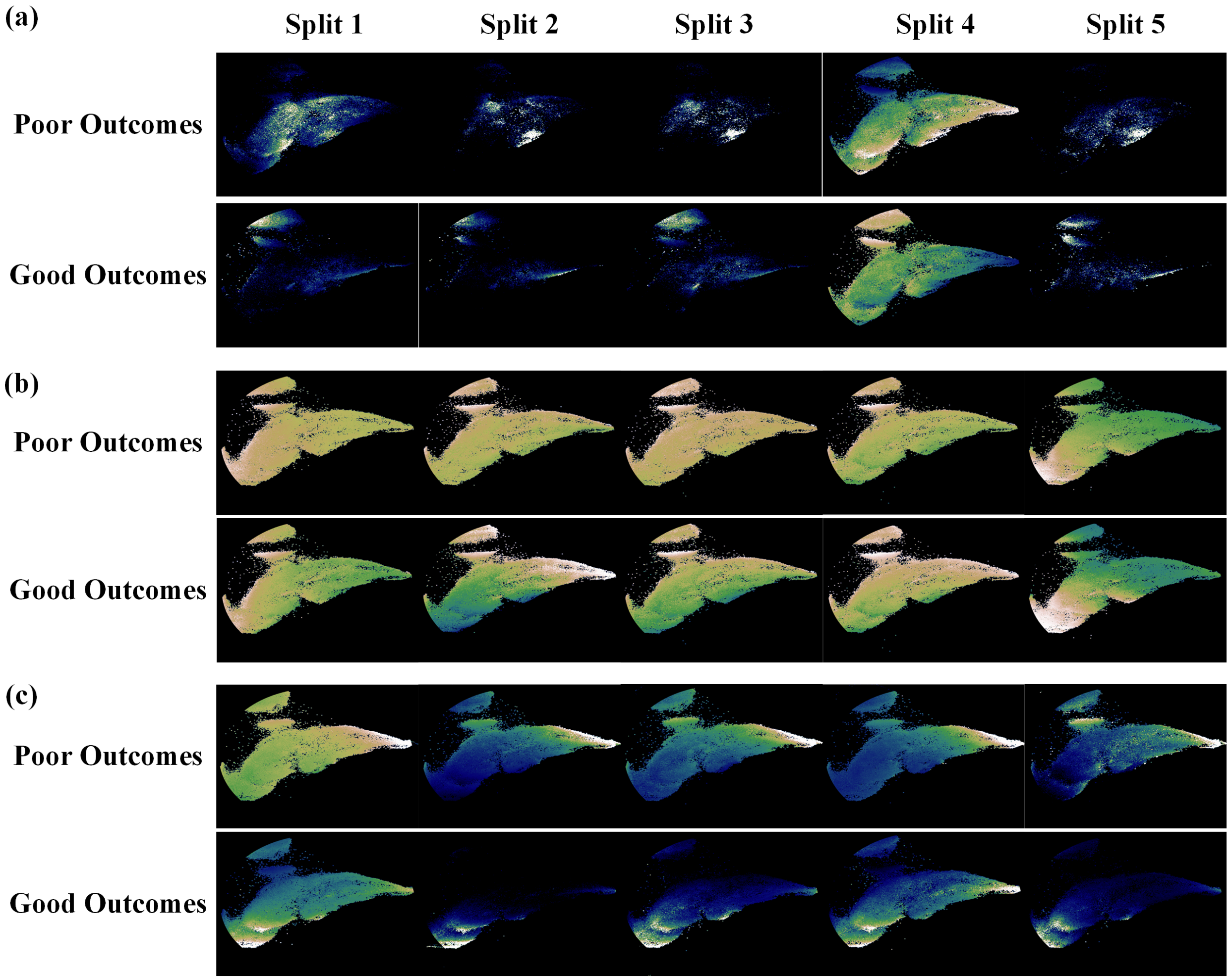


Figure 10: UMAP visualization for network attention on HIVNH dataset for (a) vMIL, (b) ind-ABMIL, and (c) dep-ABMIL.

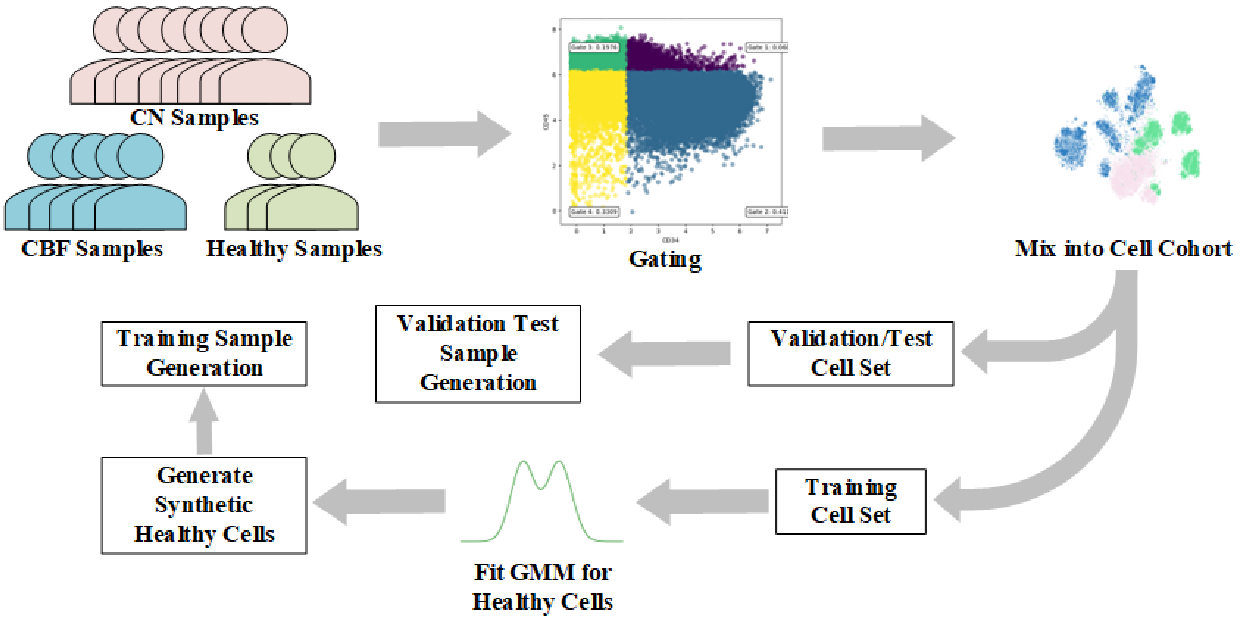


Figure 11: Dataset generation process for AML MRD task. To construct the dataset, samples are first gated to isolate blast populations. These gated blasts are pooled to create a unified cell-level cohort, which is subsequently partitioned into training, validation, and test cell sets. For the training set, a GMM is fitted using healthy cells from the training cohort, enabling the generation of synthetic healthy cells. These synthetic cells are then combined with real CN and CBF leukemic cells to produce training samples. In contrast, the validation and test sets are constructed exclusively from real cells sampled directly from their respective partitions.

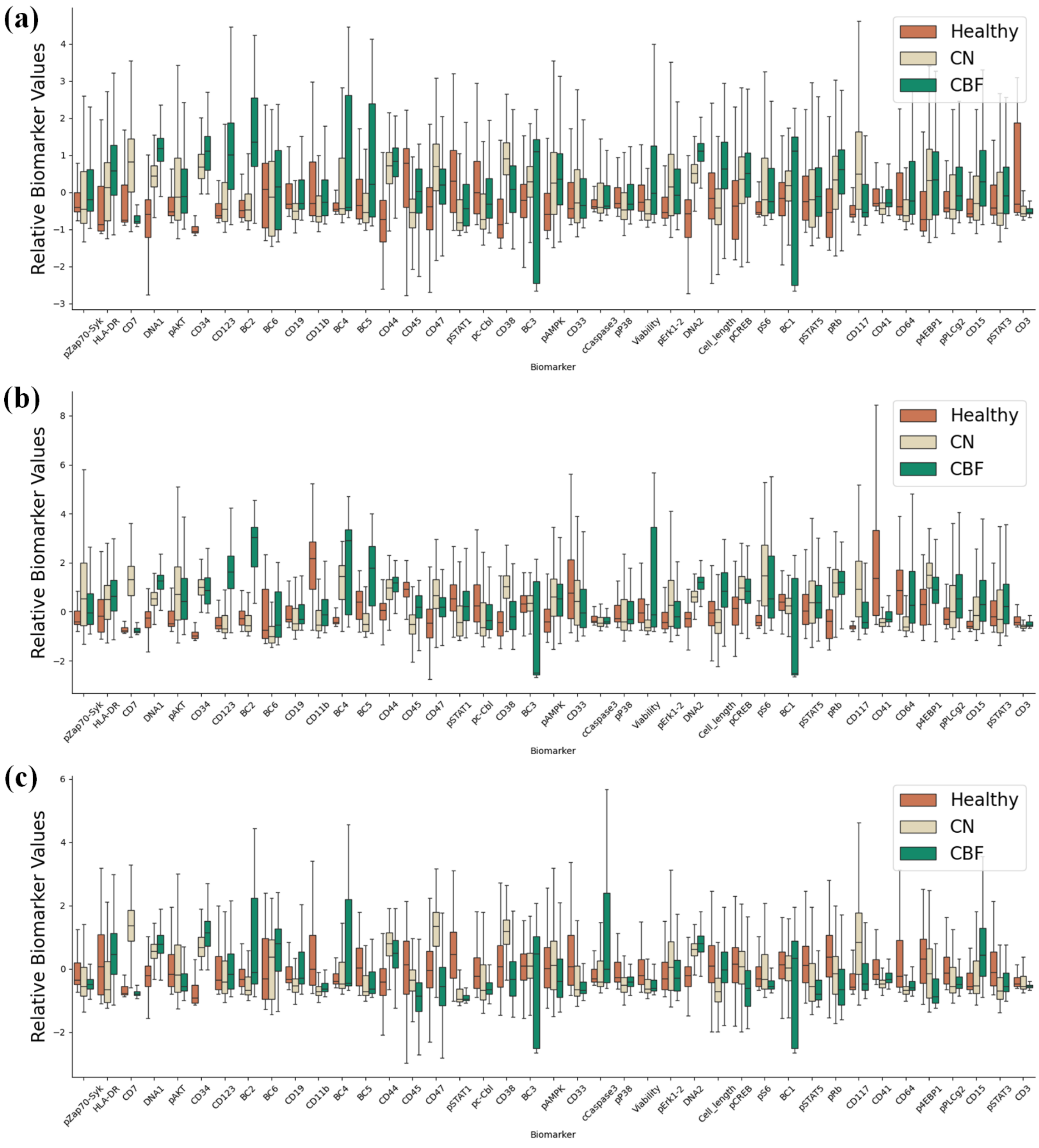


Figure 12: Biomarker values are statistically analyzed for cell populations derived from healthy, CN, and CBF-associated sampled cells. (a) Random sampling, (b) ind-ABMIL, and (c) dep-ABMIL.

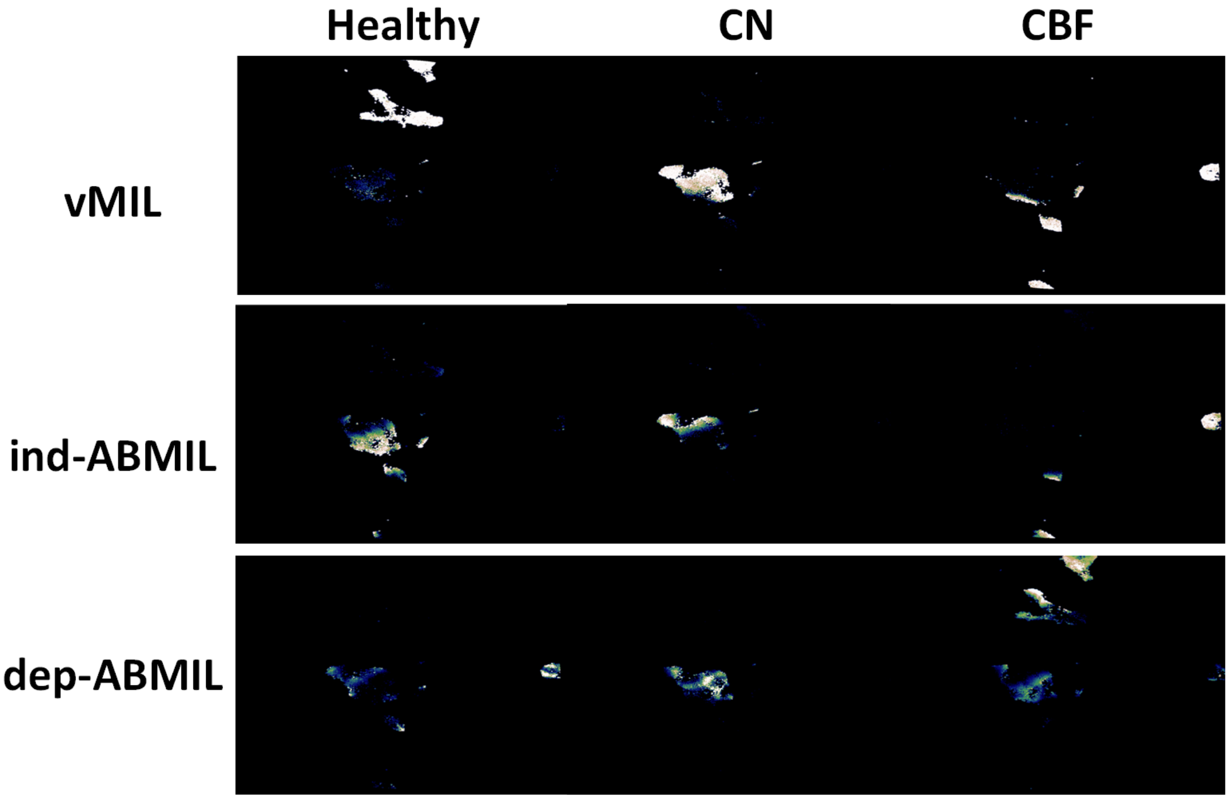


Figure 13: UMAP visualization for network attention on AML dataset for (a) vMIL, (b) ind-ABMIL, and (c) dep-ABMIL.

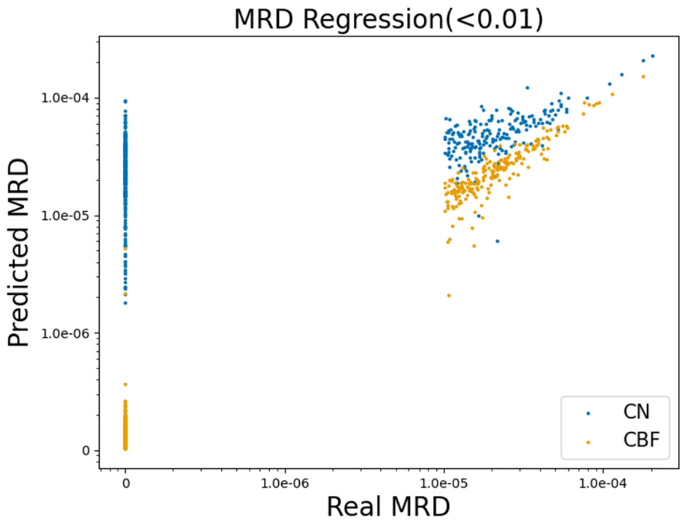


Figure 14: MRD prediction performance in challenge with maximum MRD ratio $1 \times 10^{-4}$.

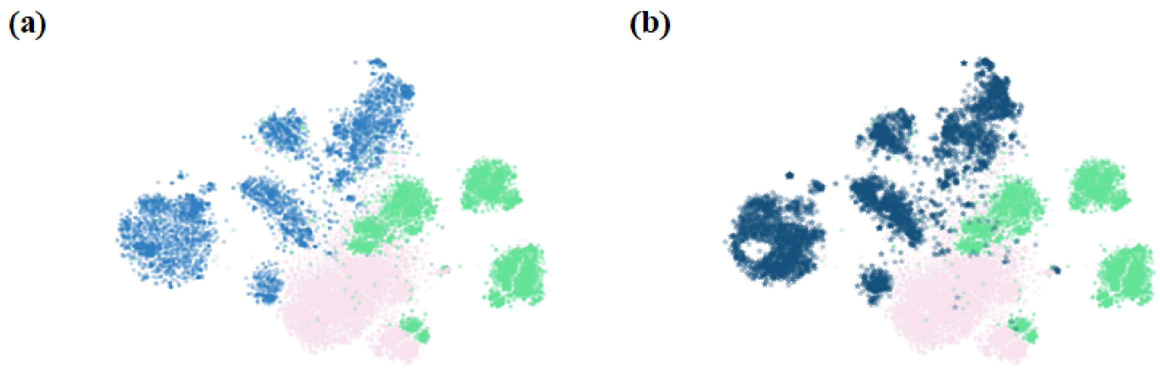


Figure 15: Visualization of UMAP space distribution for (a) real cell and (b) synthetic healthy cell distributions. Blue dot represents healthy cells while the other two are AML blasts.

(a)

(b)

(c)

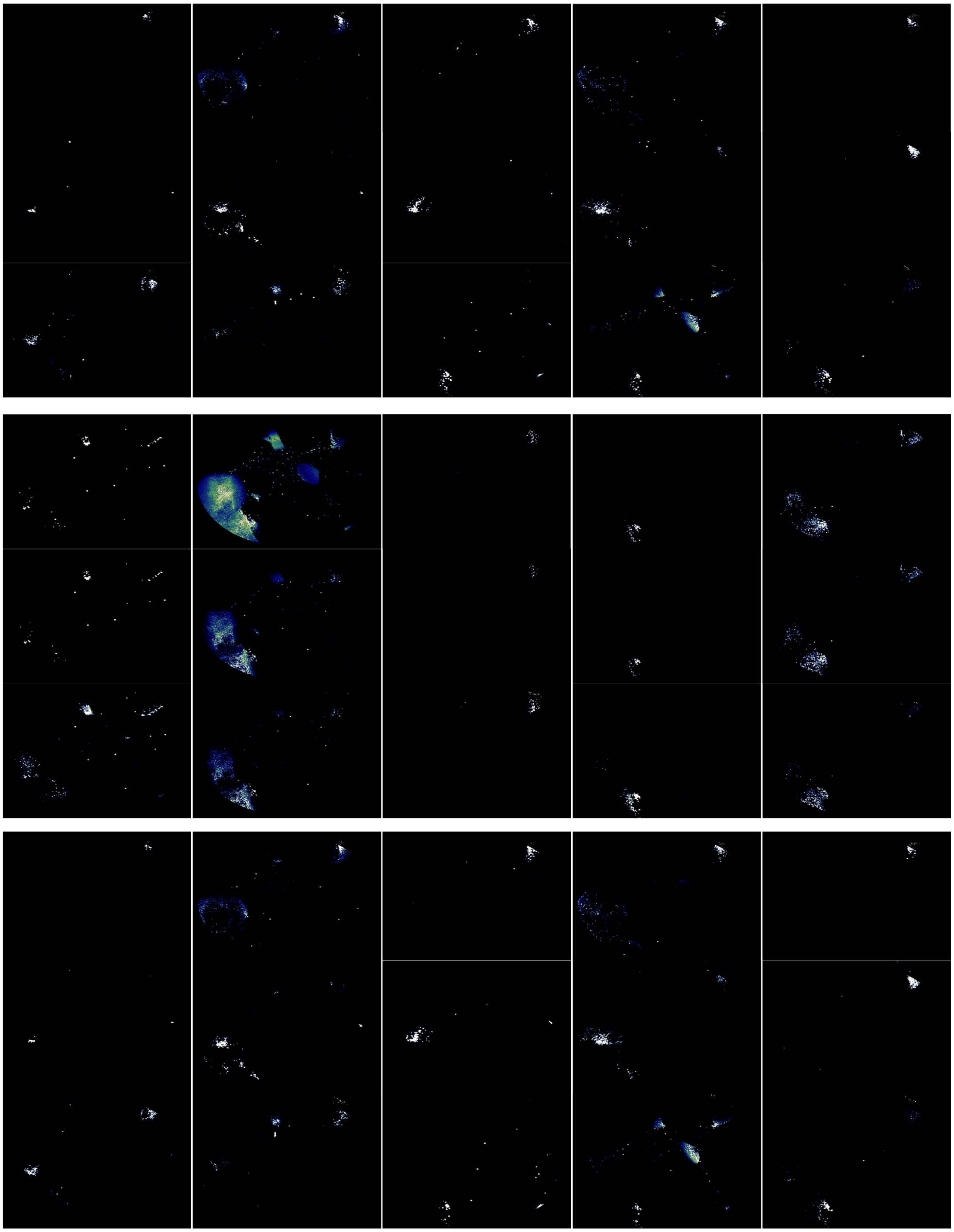

Figure 16: Learned instance attention distribution for COVID dataset panel 1 visualization of UMAP space distribution across different split (columns) for (a) single panel training, (b) multi-panel model with random initialization, and (c) multi-panel model with pretrained weight from corresponding single panel model.

(a)

(b)

(c)

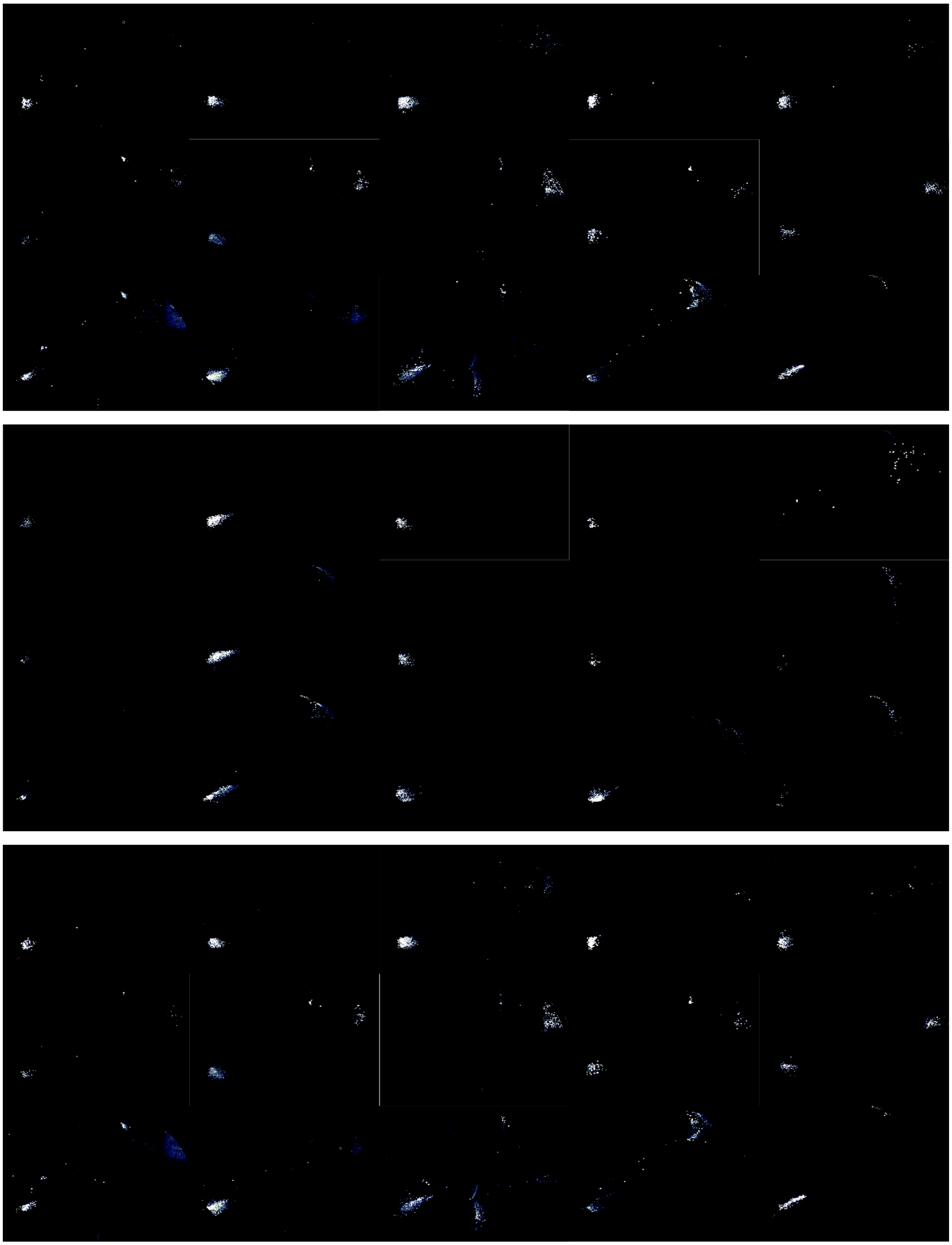

Figure 17: Learned instance attention distribution for COVID dataset panel 2 visualization of UMAP space distribution across different split (columns) for (a) single panel training, (b) multi-panel model with random initialization, and (c) multi-panel model with pretrained weight from corresponding single panel model.

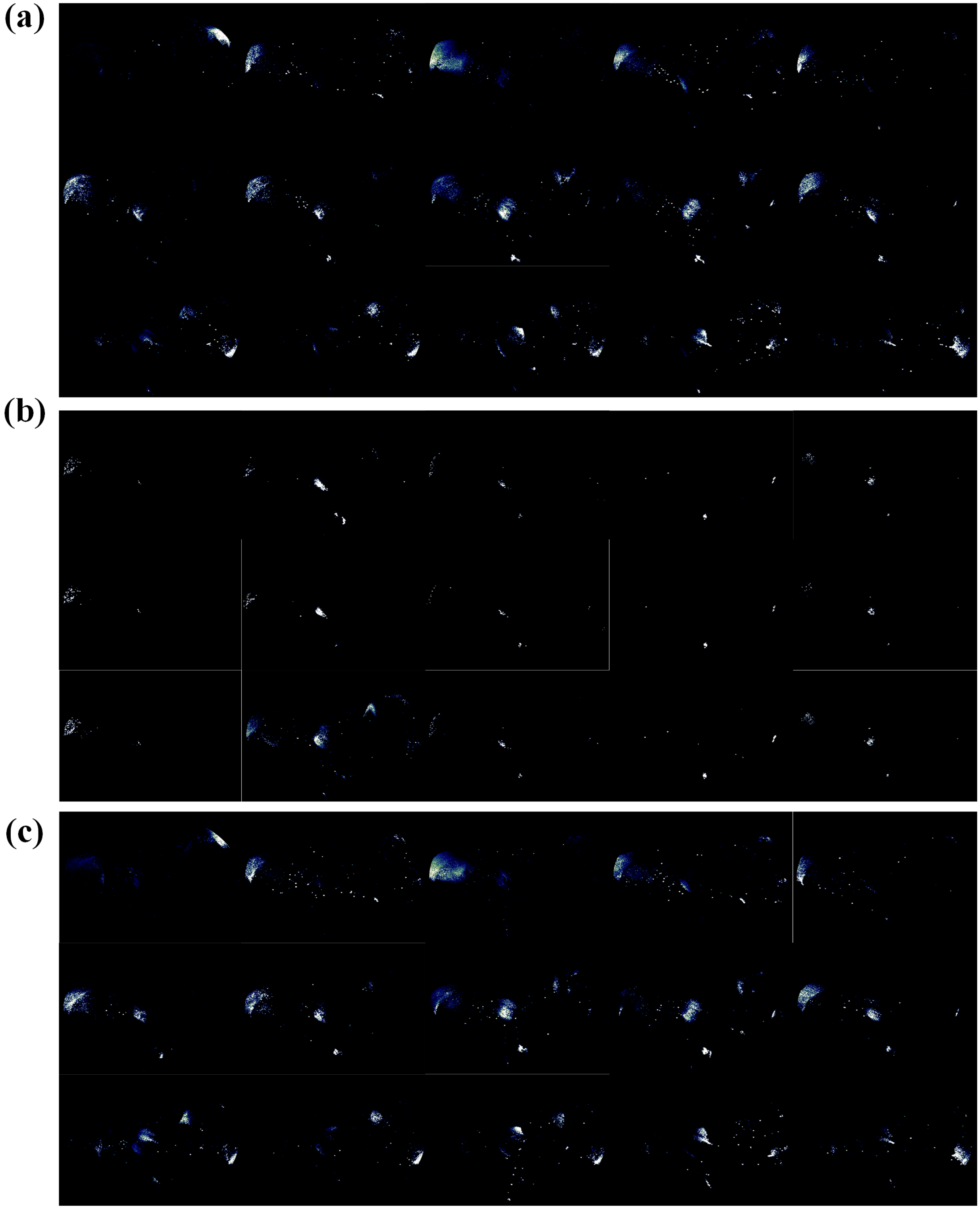

Figure 18: Learned instance attention distribution for COVID dataset panel 3 visualization of UMAP space distribution across different split (columns) for (a) single panel training, (b) multi-panel model with random initialization, and (c) multi-panel model with pretrained weight from corresponding single panel model.

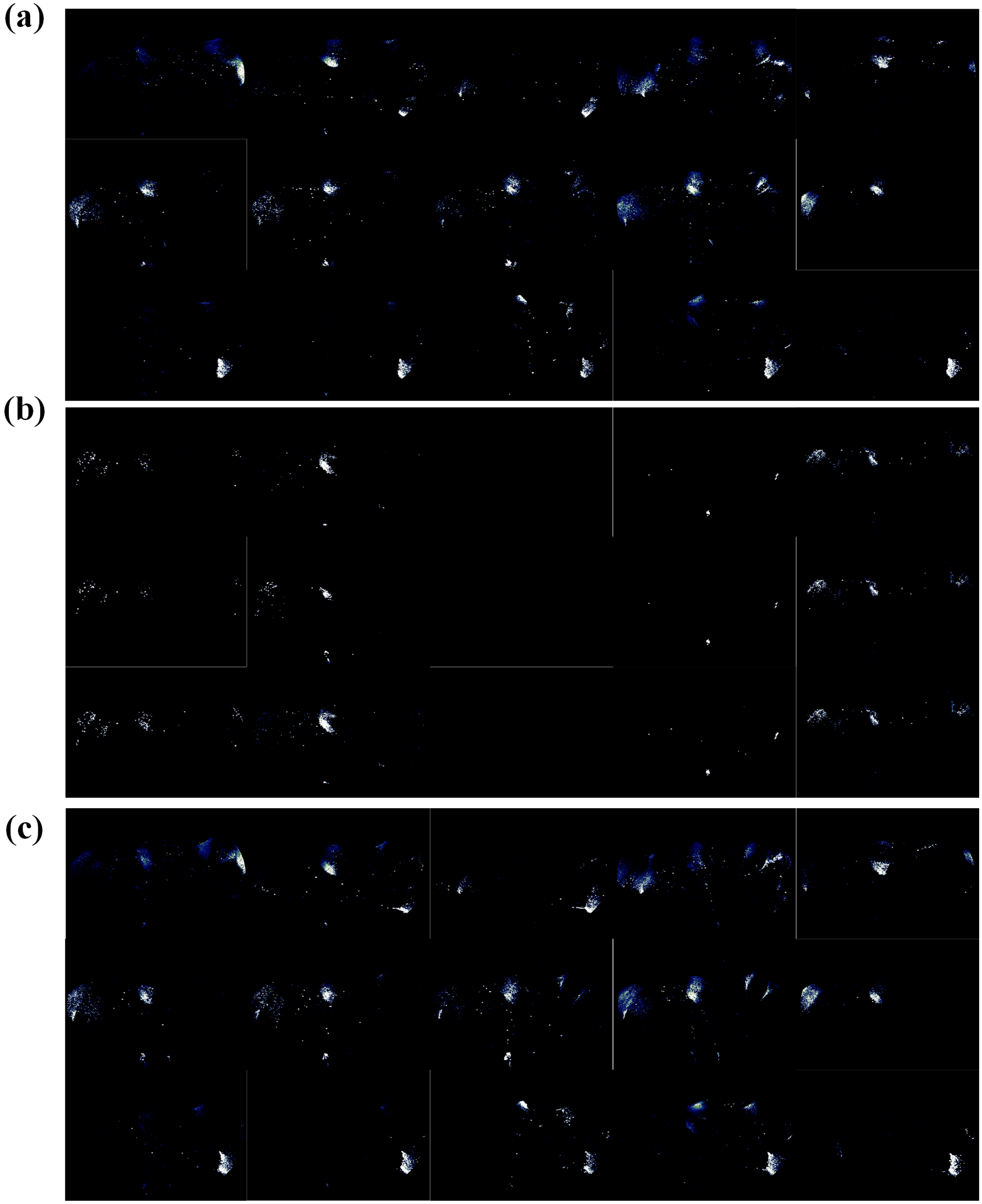


Figure 19: Learned instance attention distribution for COVID dataset panel 4 visualization of UMAP space distribution across different split (columns) for (a) single panel training, (b) multi-panel model with random initialization, and (c) multi-panel model with pretrained weight from corresponding single panel model.

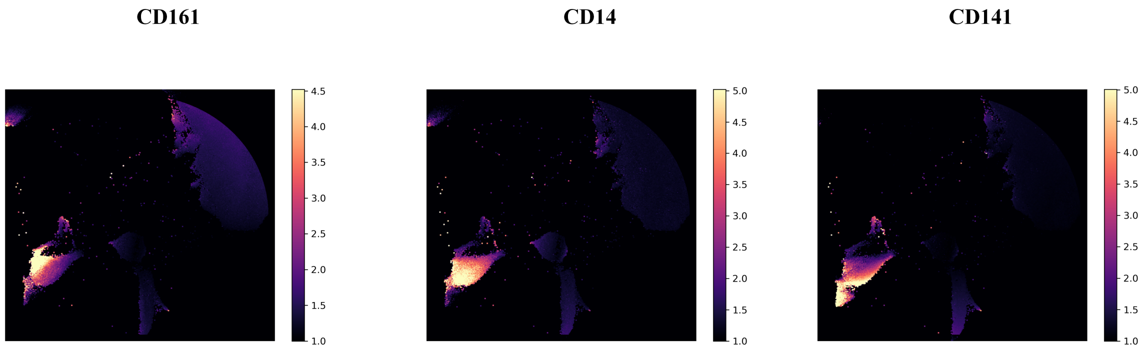


Figure 20: Selected biomarker distribution for COVID dataset panel 2 visualization of UMAP space. Compared to the phenotype distribution, we can find the healthy cell populations are associated with CD161+ while COVID severe cell populations are associated with CD14+ or CD141+ cell populations.
